# Antizyme regulates polyamine uptake via ATP13A3

**DOI:** 10.64898/2026.07.01.735194

**Authors:** Christoph Müller, Gabriela Casanova-Sepúlveda, Alessandra Blanco Hernandez, Dirk Mossmann, Markus Oppelt, Josephine K. Carscadden, Marco Colombi, Danilo Ritz, Michael N. Hall

## Abstract

Intracellular polyamine levels are tightly regulated and frequently elevated in cancer. While the regulation of polyamine synthesis is well characterized, the regulation of polyamine uptake is poorly understood. Here we identify ATP13A3 as a plasma membrane polyamine transporter. An increase in intracellular polyamine levels, due to polyamine supplementation or induction of polyamine synthesis, causes rapid internalization of the transporter and inhibition of polyamine uptake. Mechanistically, increased polyamine concentrations lead to expression of antizyme (AZ) which binds ATP13A3 and triggers its internalization. Mutations in the AZ binding site of ATP13A3 prevent AZ binding and lead to polyamine toxicity through uncontrolled polyamine influx. These findings establish ATP13A3 as a key polyamine transporter and provide a framework for targeting polyamine metabolism in cancer.

## Introduction

Polyamines, putrescine, spermidine, and spermine, are essential polycationic metabolites present at up to millimolar concentrations in cells (Fig. 1A) (*1*). Owing to their positive charge, they interact with a variety of cellular components, including proteins, lipids, and nucleic acids, and are required for proliferation and differentiation (*2, 3*). Polyamines are also implicated in disease and aging (*4, 5*). Cells produce polyamines or obtain them from the diet or the microbiome (*6*). Polyamine biosynthesis and uptake are regulated to ensure sufficient intracellular polyamines to sustain growth and proliferation (*7*). Polyamine homeostasis is also important to avoid high, toxic concentrations, particularly of spermidine and spermine (*8*). However, the coordinated regulation of synthesis and uptake that mediates polyamine homeostasis is poorly understood.

**Fig. 1.**
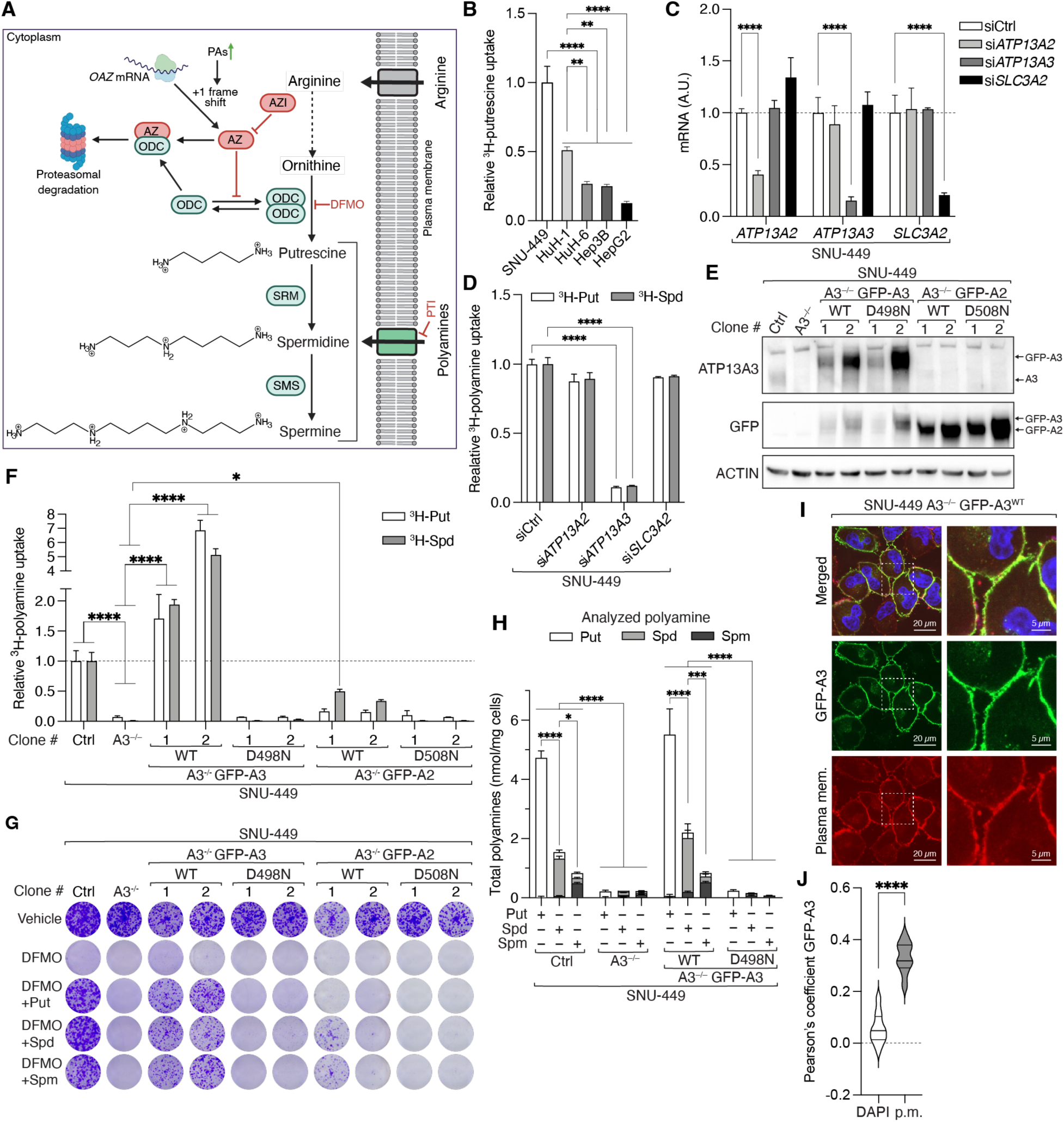
ATP13A3 is a plasma membrane polyamine transporter. **(A)** Schematic representation of polyamine metabolism and chemical structures of putrescine, spermidine, and spermine. The identity of the main plasma membrane polyamine transporter remains unknown. (**B**) Relative ^3^H-putrescine uptake into SNU-449, HuH-1, HuH-6, Hep3B, HepG2 cells. n=3. (**C**) mRNA levels of *ATP13A2*, *ATP13A3*, and *SLC3A2* in SNU-449 cells upon treatment with siCtrl, si*ATP13A2*, si*ATP13A3*, or si*SLC3A2.* n=3. (**D**) Relative ^3^H-polyamine uptake into SNU-449 cells treated with siRNAs from (C). n=3. (**E**) Immunoblots of ATP13A3 and GFP in SNU-449 control, A3^-/-^ and GFP-A3 or GFP-A2 expressing cells. ACTIN serves as loading control. (**F**) Relative ^3^H-polyamine uptake of cell lines from (E). n=4 (**G**) Representative clonogenic growth assay of cell lines from (E) treated with PBS, 1 mM DFMO, or with 1 mM DFMO and individual 1 µM polyamines. (**H**) Targeted metabolomic analysis of selected cell lines from (E) depleted of polyamines (1 mM DFMO for 2 days) and treated with 100 µM putrescine, spermidine, or spermine for 1 h. Individual polyamine contributions are stacked. n=3. (**I** and **J**) Fluorescence microscopy of DAPI, GFP-A3^WT^ and a plasma membrane marker in SNU-449 A3^-/-^ GFP-A3^WT^ cells. *p < 0.05, **p < 0.01, ****p < 0.0001 by one-way-ANOVA (B), two-way-ANOVA (C, D, F, H), or t-test (J).

Polyamine biosynthesis starts with the production of putrescine from ornithine, catalyzed by the rate-limiting enzyme ornithine decarboxylase (ODC) (Fig. 1A). Spermidine and spermine are then sequentially produced from putrescine by the addition of aminopropyl groups, catalyzed by spermidine synthase (SRM) and spermine synthase (SMS), respectively. Polyamine biosynthesis is tightly regulated through a multi-layered feedback mechanism. Increased intracellular polyamine levels lead to the expression of ODC antizyme (hereafter referred to as AZ) via a ribosomal frameshift mechanism (*9*). AZ forms a heterodimer with ODC to inhibit its activity and induce its proteasomal degradation. Low intracellular polyamine levels lead to the proteasomal degradation of AZ, thereby preventing further inhibition of ODC. In addition, antizyme inhibitor (AZI), a structural homolog of ODC that lacks enzymatic activity, binds AZ with higher affinity than ODC, thereby sequestering AZ and preventing ODC degradation (*10*).

Evidence that AZ also inhibits polyamine uptake was first reported more than 30 years ago (*11, 12*). However, the underlying molecular mechanism of uptake inhibition remains unknown, possibly because polyamine transport in metazoans is poorly characterized (*13, 14*). Recently, several polyamine transporters have been reported, including SLC3A2 and four P5-type ATPases, ATP13A2 through 5 (*15–19*). ATP13A2-5 are multi-domain transporters that bind ATP via their nucleotide-binding domain (NBD) (*18*). The binding and subsequent hydrolysis of ATP lead to conformational changes that mediate polyamine transport. Indeed, ATPase-deficient, mutant ATP13A2 (D508N) and ATP13A3 (D498N) are defective for polyamine transport (*17, 20*). ATP13A2 and A3 have been proposed to reside mainly in endolysosomal compartments where they transport endocytosed polyamines from the lumen to the cytoplasm (*21*). ATP13A3 has also been suggested to reside at the plasma membrane in a pancreatic cancer cell line under certain conditions, but whether it is active at the plasma membrane remains unclear (*22*).

## Results

### ATP13A3 is a plasma membrane polyamine transporter

We recently showed that murine and human hepatocellular carcinoma (HCC) present high levels of polyamines despite downregulated arginine-to-polyamine conversion (*23*). Liver tumors achieve elevated polyamine concentrations by upregulating polyamine import. To study further polyamine import, we sought to identify cells with particularly high polyamine uptake. We assayed polyamine transport in a panel of HCC cells of which SNU-449 cells showed the highest basal ^3^H-putrescine uptake (Fig. 1B). Inhibition of polyamine biosynthesis in SNU-449 cells, by treatment with the ODC inhibitor DFMO, prevented cell growth unless the cells were supplemented with exogenous polyamines (fig. S1A). The ability of exogenous polyamines to sustain growth in this context was abrogated by co-treatment with the polyamine transport inhibitor (PTI) AMXT-1501. We thus focused on SNU-449 cells as a valid system to characterize polyamine transport. Several proteins have been suggested to mediate polyamine uptake, among which ATP13A2, ATP13A3 and SLC3A2 are expressed in SNU-449 cells (*16, 17, 24*). Knockdown exclusively of ATP13A3 decreased ^3^H-putrescine and ^3^H-spermidine uptake (Fig. 1, C and D, fig. S1, B and C), indicating that ATP13A3 is a particularly important polyamine transporter.

Knockout (KO) of ATP13A3 (A3^-/-^) in SNU-449 cells reduced putrescine and spermidine uptake by >90% and abolished clonogenic growth upon inhibition of polyamine synthesis (fig. S1, D to F). Confirming our knockdown results, ATP13A2 KO (A2^-/-^) had no effect on polyamine uptake or clonogenic growth (fig. S1, G to J). Expression of N-terminally GFP-tagged, wildtype ATP13A3 (GFP-A3^WT^) in A3^-/-^ cells restored polyamine uptake and growth, whereas expression of a GFP-tagged, ATPase-deficient mutant ATP13A3 D498N (GFP-A3^D498N^) did not restore uptake (Fig. 1, E to G). To investigate the substrate selectivity of ATP13A3, we assayed intracellular accumulation of specific polyamines, by liquid chromatography coupled to mass spectrometry (LC-MS), in SNU-449 cells containing or lacking ATP13A3 (Fig. 1H). All three polyamines accumulated in an ATP13A3-dependent manner, although putrescine accumulated at a faster rate than spermidine or spermine. Thus, ATP13A3 is a broad polyamine transporter with a possible preference for putrescine over spermidine and spermine. We note that ATP13A2 was reported to show reverse selectivity, preferring spermidine and spermine over putrescine (*17*). Interestingly, GFP-tagged ATP13A2 WT (GFP-A2^WT^) only weakly complemented an ATP13A3 deficiency despite being highly expressed (Fig. 1, E to G). The complementation by ATP13A2 was only observed for spermidine and spermine supplementation, in line with its reported selectivity (Fig. 1, F and G).

To examine the subcellular localization of ATP13A3, we performed fluorescence microscopy on SNU-449 A3^-/-^ cells expressing GFP-A3^WT^. These cells exhibited clear GFP-A3^WT^ localization at the plasma membrane (Fig. 1, I and J), contrary to the transporter’s previously proposed localization in endolysosomal compartments (*21*). To investigate further the plasma membrane localization, we expressed GFP-A3^WT^ in HepG2 and HeLa cell lines. These cells displayed intracellular GFP-A3^WT^ with little-to-no plasma membrane localization. However, upon inhibition of polyamine synthesis with DFMO, intracellular GFP-A3^WT^ translocated to the plasma membrane (fig. S2, A to E). In contrast, GFP-A3^WT^ in SNU-449 cells exhibited strong plasma membrane localization independent of DFMO treatment (fig. S2, A, F, and G). Unlike GFP-A3^WT^, GFP-A2^WT^ in SNU-449, HeLa and HepG2 cells, exhibited intracellular localization that correlated with LAMP1 staining and did not change upon DFMO treatment (fig. S2, A and H to M). Overall, our findings suggest ATP13A3 is a plasma membrane transporter. ATP13A2, as reported previously, exhibits intracellular endolysosomal localization (*17*). The differential localization of the two transporters suggests they perform distinct physiological functions and thus provides an explanation for the inability of ATP13A2 to complement an ATP13A3 deficiency.

**Fig. 2.**
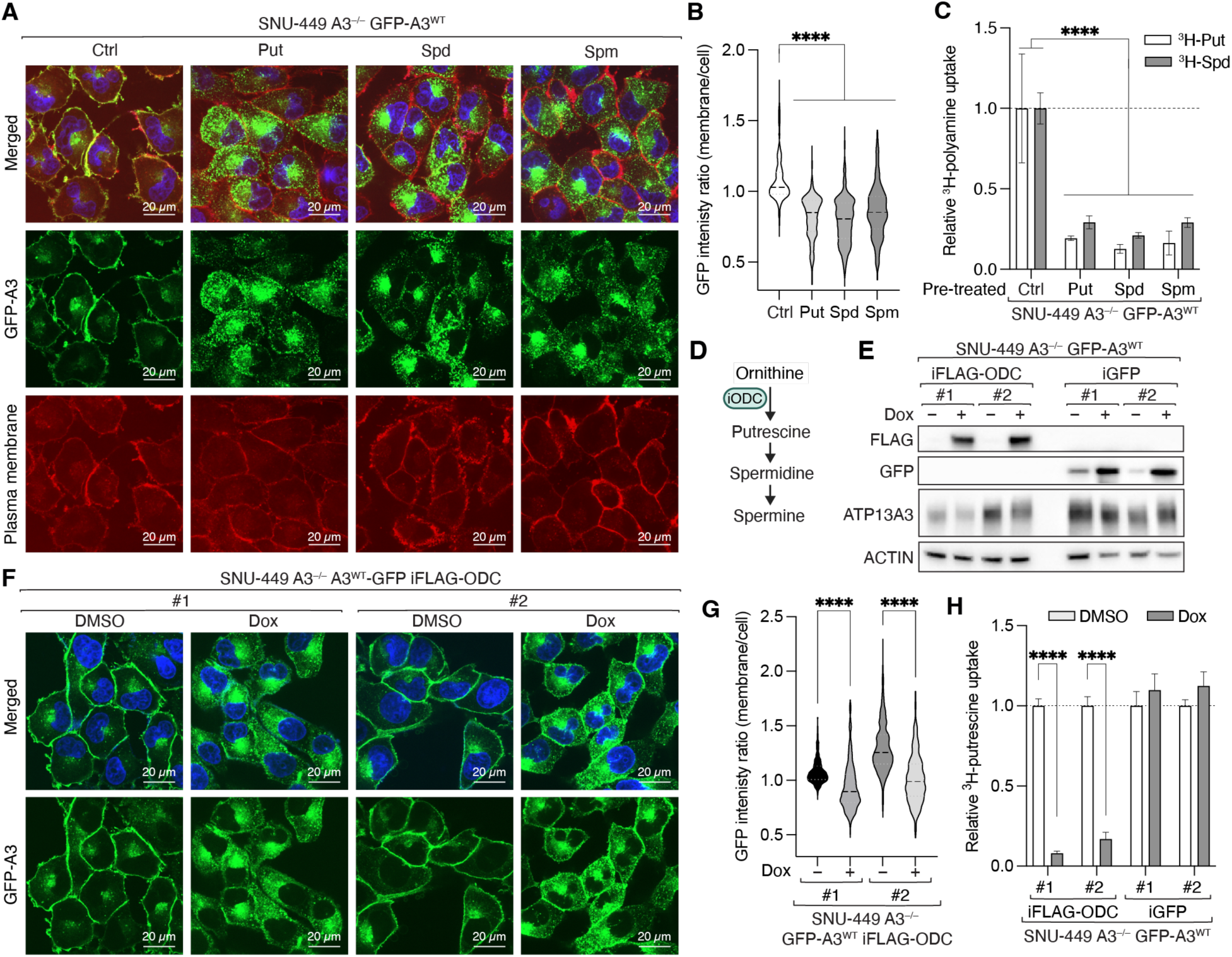
ATP13A3 is internalized upon increasing polyamine levels. **(A** and **B)** Fluorescence microscopy of SNU-449 A3^-/-^ GFP-A3^WT^ cells treated with PBS or 10 µM polyamines for 2 h. (**C**) Relative ^3^H-polyamine uptake into SNU-449 A3^-/-^ GFP-A3^WT^ cells treated with PBS or 10 µM polyamines for 2 h. n=4. (**D**) Schematic representation of inducible FLAG-ODC (iFLAG-ODC) expression. (**E**) Immunoblots of FLAG, GFP, and ATP13A3 in control or doxycycline-induced SNU-449 A3^-/-^ GFP-A3^WT^ iFLAG-ODC or iGFP cells. ACTIN serves as loading control. (**F** and **G**) Fluorescence microscopy of DAPI and GFP-A3^WT^ in SNU-449 A3^-/-^ GFP-A3^WT^ iFLAG-ODC cells treated with DMSO or doxycycline. (**H**) Relative ^3^H-putrescine uptake into cells from (E). n=4. ****p < 0.0001 by one-way-ANOVA (B) or two-way-ANOVA (C, G, H).

### ATP13A3 is internalized upon increased intracellular polyamine levels

Unlike GFP-A3^WT^ expressing HeLa and HepG2 cell lines, GFP-A3^WT^ expressing SNU-449 cells exhibited clear plasma membrane localization even in the absence of DFMO (see above). Given that SNU-449 cells have particularly low polyamine biosynthesis (*23*), we reasoned that low intracellular polyamine availability drives constitutive plasma membrane targeting of ATP13A3 in these cells. To test whether ATP13A3 localization is regulated by intracellular polyamine levels, we treated SNU-449 A3^-/-^ GFP-A3^WT^ cells with polyamines. We observed nearly complete internalization of the transporter within 2 h, regardless of the polyamine added (Fig. 2, A and B). To assess if internalization was specific to polyamines, we also treated cells with ornithine or arginine, neither of which influenced GFP-A3^WT^ localization (fig. S3, A and B). A3^-/-^ GFP-A3^WT^ and SNU-449 control cells pre-treated with polyamines showed a strong reduction in polyamine uptake, suggesting that polyamine uptake requires ATP13A3 at the plasma membrane (Fig. 2C; fig. S3C). In addition, internalization requires transporter activity as ATPase-deficient GFP-A3^D498N^ remained at the plasma membrane upon putrescine treatment (fig. S3, D and E). Internalized ATP13A3 partly colocalized with the early endosome marker EEA1 and the Golgi-resident protein GM130 (fig. S3, F to I) as reported previously (*16, 21*). GFP-A3^WT^ internalization upon putrescine treatment was also observed in HeLa and HepG2 cells (fig. S3, J to M).

The above findings suggest two possibilities for the regulation of polyamine-induced ATP13A3 internalization. First, cell surface ATP13A3 binds external polyamines which then induces or signals internalization of the transporter. Second, the transporter responds to intracellular polyamine levels. In either case, transporter internalization could be a mechanism to prevent further uptake of polyamines, thereby protecting cells from accumulating high, toxic polyamine levels. To test these models, we stimulated polyamine biosynthesis by inducing expression of FLAG-tagged ODC (iFLAG-ODC) (Fig. 2, D and E). Like the external addition of polyamines, enhanced ODC expression was sufficient to induce internalization of GFP-A3^WT^ and a corresponding reduction in polyamine uptake (Fig. 2, F to H). Thus, ATP13A3 localization and thereby polyamine uptake is regulated in response to intracellular polyamine levels.

### AZ binds and promotes internalization of ATP13A3

To investigate the mechanism of ATP13A3 internalization, we performed GFP-trap pull-down experiments using SNU-449 A3^-/-^ GFP-A3^WT^ cells grown under conditions where GFP-A3^WT^ is at the plasma membrane (Ctrl) or internalized (putrescine-treated). Among the few GFP-A3^WT^ binding proteins enriched after putrescine treatment was the ODC regulator AZ1 (Fig. 3A). AZ1 is one of three AZ paralogs (AZ1-3) (*10*). AZ1 and 2 are ubiquitously expressed, although AZ2 at much lower levels, while AZ3 is restricted to the testis (*10, 25*).

**Fig. 3.**
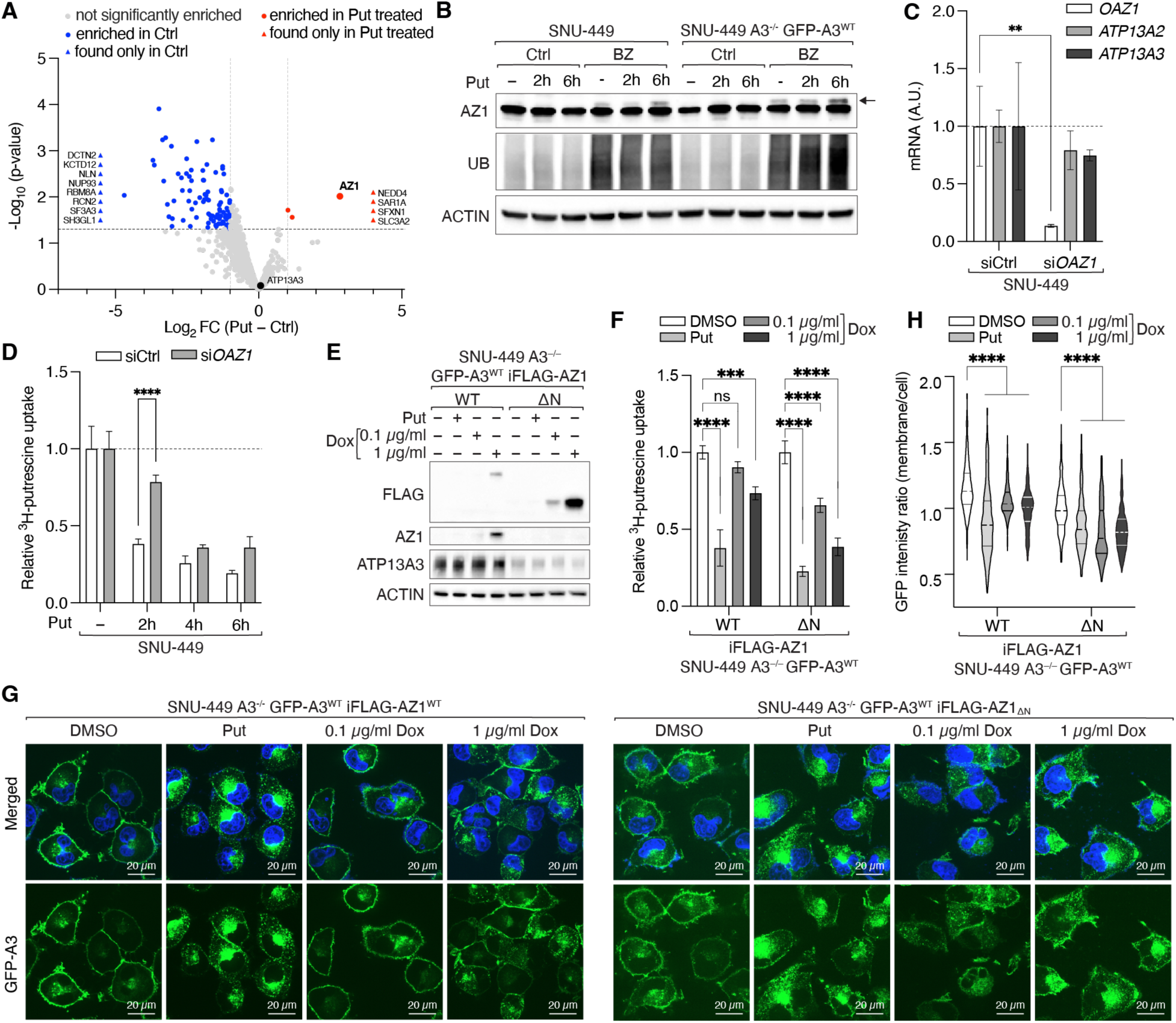
Antizyme expression leads to internalization of ATP13A3. **(A)** Volcano plot of the log_10_ (p value) against the log_2_ fold-change of proteins identified by MS after anti-GFP pull-down from putrescine-treated vs. untreated SNU-449 A3^-/-^ GFP-A3^WT^ cells. Significantly enriched (circles) or uniquely identified (triangles) proteins are shown in blue and red (log_10_(p value) > 1.3, log_2_ fold-changes +/–1). (**B**) Immunoblots of ubiquitin (UB) and AZ1 in SNU-449 control or A3^-/-^ GFP-A3^WT^ cells treated with 10 µM putrescine for 0, 2 or 6 h with or without proteasomal inhibitor bortezomib (BZ). ACTIN serves as loading control. (**C**) mRNA levels of *OAZ1*, *ATP13A2*, and *ATP13A3* in SNU-449 cells treated with siCtrl or si*OAZ1*. (**D**) Relative ^3^H-putrescine uptake into SNU-449 cells treated with siCtrl or si*OAZ1* and with 10 µM putrescine for 2, 4 or 6 h prior to uptake assay. n=3. (**E**) Immunoblots of ATP13A3, FLAG, and AZ1 in SNU-449 A3^-/-^ GFP-A3^WT^ iFLAG-AZ1^WT^ or iFLAG-AZ1_ΔN_ cells treated with DMSO, 10 µM putrescine for 2 h, or doxycycline for 30 h. ACTIN serves as loading control. (**F**) Relative ^3^H-putrescine uptake into cells from (E) pre-treated as in (E). n=4 (**G** and **H**) Fluorescence microscopy of GFP-A3^WT^ in cells from (E) treated with DMSO, 10 µM putrescine for 2 h, or doxycycline. **p < 0.01, ***p < 0.001, ****p < 0.0001 by two-way-ANOVA (C, D, F, H).

As described above, AZ is expressed upon increased intracellular polyamine concentrations via a translational frameshift mechanism and, in addition to its effect on polyamine biosynthesis, has been described to inhibit polyamine uptake through an unknown mechanism (*9, 26*). We confirmed induction of AZ1 expression by putrescine in SNU-449 control and A3^-/-^ GFP-A3^WT^ cells (Fig. 3B). Of note, owing to the short half-life of AZ1 (*27*), expression was detectable by immunoblotting only in cells treated with the proteasomal inhibitor bortezomib (BZ). Knockdown of AZ1 attenuated the reduction in polyamine uptake in response to 2 h putrescine treatment (Fig. 3, C and D). Inducible overexpression of FLAG-tagged full-length (WT) or an N-terminally truncated version (AZ1ΔN, residues 1-90 deleted) of AZ1 (iFLAG-AZ1) resulted in internalization of GFP-A3^WT^ and decreased polyamine uptake (Fig. 3, E to H). Thus, the polyamine sensor AZ1 binds and promotes internalization of ATP13A3. We note that the normally disordered N-terminus of AZ1 was deleted in AZ1ΔN to eliminate a degron and stabilize the protein.

### Identification of the ATP13A3-AZ interface

We modeled the direct binding of AZ to ATP13A3 using AlphaFold 3 (*28*). Full-length ATP13A3 and AZ1 gave a poor interface score (ipTM<0.6), likely explaining why this complex was not detected in previous *in silico* screens (fig. S4A). However, analysis of the structure of the predicted complex revealed a flexible AZ1 N-terminus that interacts with the transmembrane region of ATP13A3. This binding mode, likely non-physiological as the AZ1 N-terminus would not normally access the membrane-embedded region of the transporter, probably accounts for the low-confidence interface scores. Indeed, AZ1ΔN yielded a high-confidence interface score that predicts complex formation (Fig. 4A, ipTM=0.8). AZ1ΔN binds mostly to the NBD (residues 507-724) of ATP13A3, where it wedges into the transporter structure, creating an extensive binding interface (Fig. 4B, fig. S4B). AlphaFold 3 also predicted formation of a complex of ATP13A3 with AZ2ΔN or AZ3ΔN (fig. S4C).

**Fig. 4.**
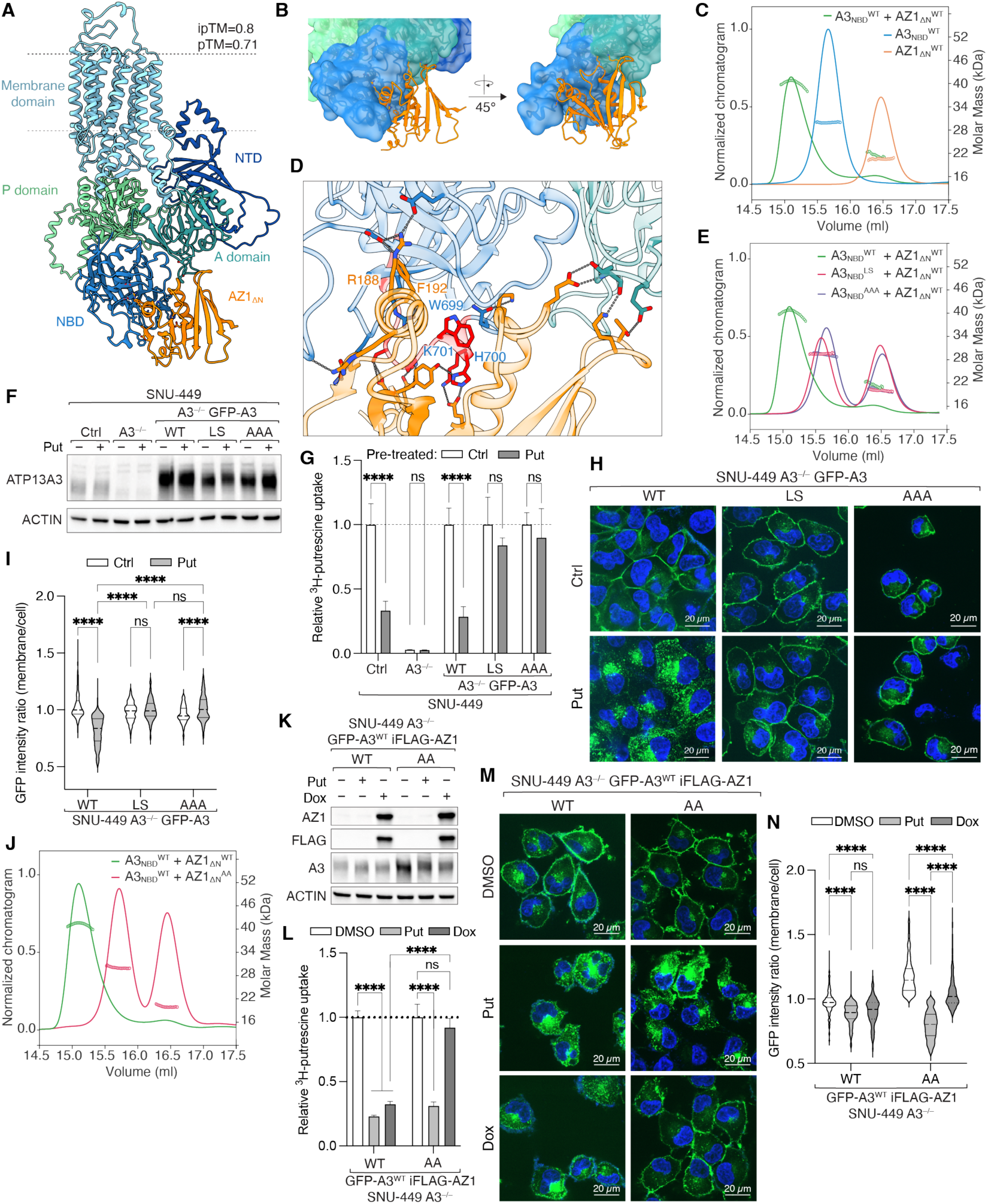
Binding of AZ1 to ATP13A3 is required for internalization. (**A**) Alphafold 3 prediction of the ATP13A3-AZ1_ΔN_ complex structure. (**B**) Surface representation of the ATP13A3 binding site. (**C**) SEC-MALS analysis of three individual injections: A3_NBD_^WT^, AZ1_ΔN_, and A3_NBD_^WT^ + AZ1_ΔN_. (**D**) Close-up of the ATP13A3-AZ1_ΔN_ interaction site. Loop swap region of ATP13A3 is highlighted in red. **(E**) SEC-MALS analysis as in (C) for: A3_NBD_^WT^ + AZ1_ΔN_, A3_NBD_^LS^ + AZ1_ΔN_, and A3_NBD_^AAA^ + AZ1_ΔN_. (**F)** Immunoblots of ATP13A3, in SNU-449 control, A3^-/-^, A3^-/-^ GFP-A3^WT^, A3^-/-^ GFP-A3^LS^ and A3^-/-^ GFP-A3^AAA^ treated with 10 µM putrescine for 2 h. ACTIN serves as loading control. (**G**) Relative ^3^H-putrescine uptake into cells from (F) pre-treated with 10 µM putrescine for 2 h. n=4. (**H** and **I**) Fluorescence microscopy of DAPI and GFP-A3 in SNU-449 A3^-/-^ GFP-A3^WT^, GFP-A3^LS^, or GFP-A3^AAA^ cells treated with 10 µM putrescine for 2 h. (**J**) SEC-MALS analysis as in (C) for A3_NBD_^WT^ + AZ1_ΔN_ and A3_NBD_^WT^ + AZ1_ΔN_^AA^. (**K**) Immunoblots of AZ1, FLAG, and ATP13A3 in SNU-449 A3^-/-^ GFP-A3^WT^ iAZ1^WT^ or iAZ1^AA^ cells treated with DMSO, 10 µM putrescine for 2 h or 1 µg/ml doxycycline for 30 h. ACTIN serves as loading control. **(L)** Relative ^3^H-putrescine uptake into cells from (K) and pre-treated as in (K). n=4. (**M** and **N**) Fluorescence microscopy of cells from (K) and treated as in (K). ****p < 0.0001 by two-way-ANOVA (G, I, L, N).

To confirm binding, we purified recombinant human ATP13A3 NBD (A3NBD) and AZ1ΔN from *E. coli* (fig. S4D). Size exclusion chromatography coupled to multi angle light scattering (SEC-MALS) analysis of the two proteins individually revealed symmetric peaks with the predicted molecular weights. Equimolar combination of the two proteins resulted in a single peak with a higher molecular weight indicating complex formation (Fig. 4C). Of note, AlphaFold 3 did not predict complex formation for any combination of ATP13A2, ATP13A4, or ATP13A5 with AZ1ΔN, AZ2ΔN, or AZ3ΔN (ipTMs<0.6, fig. S4, C and E). ATP13A3 and ATP13A2 share a high degree of sequence similarity (55%) and structural homology (RMSD = 6.544 Å); however, superimposition of the structures revealed a lower degree of conservation for the NBDs (fig. S4F). To identify the region important for AZ binding, we generated different chimeric input sequences, in which we replaced parts of the ATP13A3 NBD with the corresponding parts of ATP13A2. Swapping only residues 690-702 of ATP13A3 with 692-704 of ATP13A2 (fig. S4G) strongly decreased binding affinity of AZ1 as determined by AlphaFold 3 and Rosetta Interface Analyzer (RIA) (fig. S4H) (*29*). As the swapped sequence is part of a loop structure in ATP13A3, we named the resulting chimera A3 loop swap (A3^LS^). Analyzing the binding of the loop region to AZ1ΔN, we observed that W699, H700, K701 (referred to as the WHK motif) are wedged between two helices of AZ1 and thus likely important for the interaction with AZ1 (Fig. 4D). Purified, recombinant A3NBD^LS^ and the A3 W699A/H700A/K701A triple mutant (A3NBD^AAA^) (fig. S4D) failed to form a complex with AZ1ΔN, as determined by SEC-MALS (Fig. 4E). Full-length GFP-A3^LS^ and GFP-A3^AAA^ versions of ATP13A3 expressed in SNU-449 A3^-/-^cells were functional for polyamine uptake (fig. S4I). However, in both cases, cells were unable to decrease uptake upon putrescine treatment (Fig. 4G), consistent with the transporters remaining at the plasma membrane (Fig. 4, H and I).

AZ1 engages ATP13A3 and ODC via a similar binding surface, involving multiple contacts (fig. S4, J and K) (*30*). To identify residues in AZ1 important for ATP13A3 binding, we performed an *in silico* alanine screening of all residues within 8 Å of the predicted ATP13A3-AZ1ΔN interface using RIA (fig. S4L). This identified AZ1 R188 and F192 (referred to as a RF motif) as important for transporter binding. Purified, recombinant R188A/F192A double mutant (AZ1ΔN^AA^) failed to bind A3NBD^WT^ as assessed by SEC-MALS (Fig. 4J). Furthermore, induction of iAZ^AA^ failed to alter the plasma membrane localization of GFP-A3^WT^ or perturb polyamine uptake, whereas induction of endogenous AZ1 by putrescine treatment triggered GFP-A3^WT^ internalization. (Fig. 4, K to N). Thus, AZ1 and ATP13A3 interact via specific sequences to mediate polyamine-induced internalization of the transporter.

### AZ-mediated ATP13A3 regulation is physiologically relevant

High intracellular polyamine levels are toxic, whereas low levels are unable to sustain cell proliferation and function. Does altered expression of AZ affect cell growth? In other words, is regulation of polyamine transport by AZ physiologically relevant? To answer this question, we first examined whether sustained expression of AZ prevents growth of cells that rely on exogenous polyamine. Indeed, clonogenic growth of SNU-449 cells expressing AZ1 and supplemented with exogenous polyamines was decreased upon treatment with DFMO (Fig. 5A, fig. S5A). This effect was more pronounced upon spermidine or spermine supplementation, likely due to their lower uptake rates compared to putrescine (see above). Second, we examined whether loss of AZ causes polyamine toxicity. Induced AZ1 knockdown in SNU-449 A3^-/-^ GFP-A3^WT^ cells increased spermidine and spermine toxicity independent of effects on polyamine synthesis (fig. S5, B and C). Spermidine and spermine are generally more cytotoxic than putrescine, in part because their catabolism by polyamine oxidases generates reactive oxygen species and toxic aldehydes such as acrolein (*31*). Finally, we assessed spermidine toxicity in cells expressing ATP13A3 variants or lacking ATP13A3 (Fig. 5, B and C). While SNU-449 control cells expressing endogenous ATP13A3 or A3^-/-^ cells expressing recombinant GFP-A3^WT^ were sensitive to high levels of exogenous spermidine, A3^-/-^ cells were completely insensitive to spermidine. Strikingly, A3^-/-^ cells expressing GFP-A3^LS^, the ATP13A3 variant that is unable to bind AZ, were highly sensitive to spermidine. Thus, AZ-mediated down-regulation of ATP13A3 is physiologically relevant in mediating polyamine homeostasis.

**Fig. 5.**
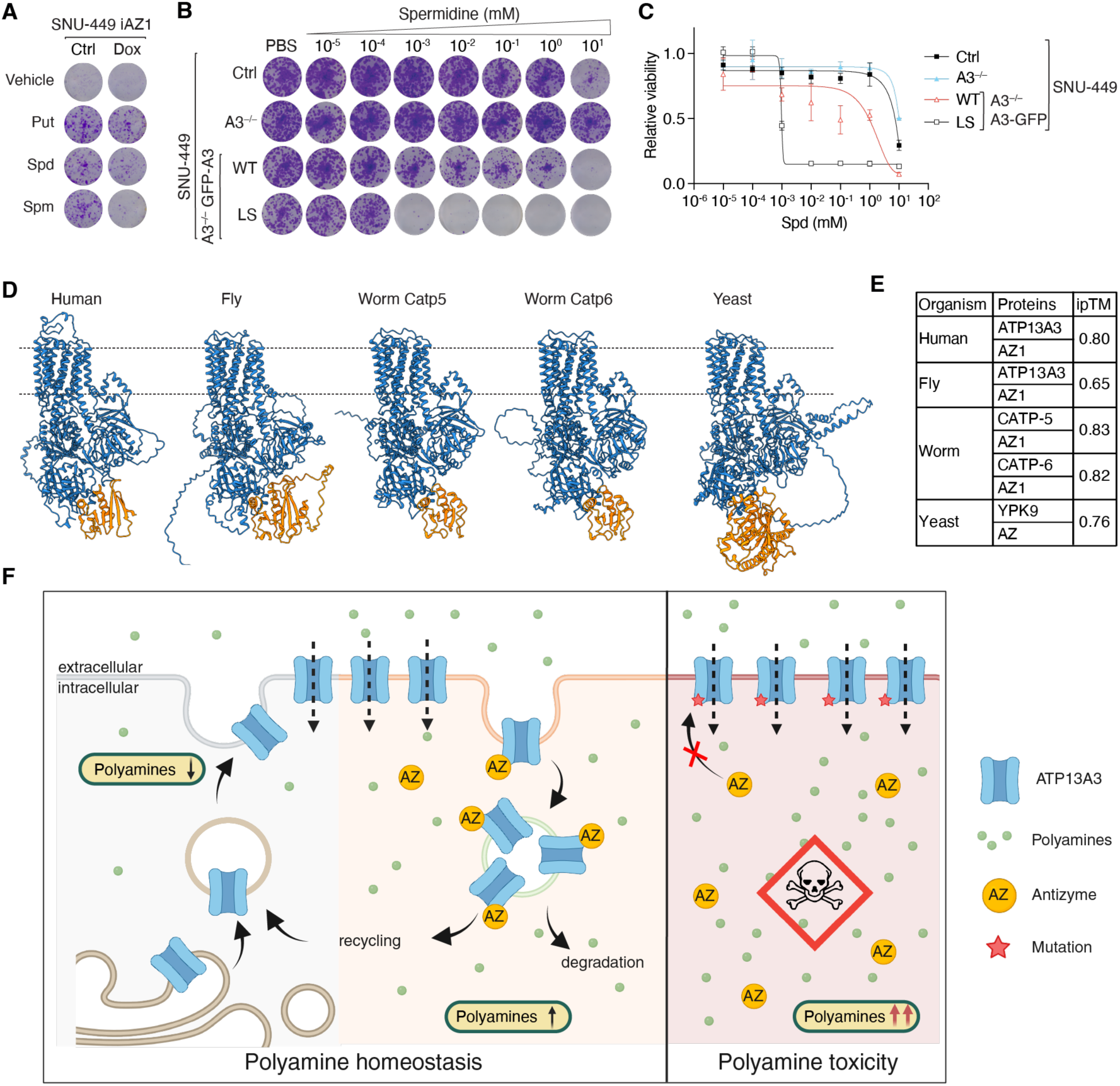
Conservation of the ATP13A3-AZ1 interaction across eukaryotes confers protection against polyamine toxicity. **(A)** Representative clonogenic growth assay of SNU-449 A3^-/-^ GFP-A3 iAZ1 cells treated with 1 mM DFMO, 1 µg/ml doxycycline, and individual 1 µM polyamines. (**B**) Representative clonogenic growth assay of SNU-449 control, A3^-/-^, A3^-/-^ GFP-3^WT^, A3^-/-^ GFP-A3^LS^, and A3^-/-^ GFP-A3^AAA^ cells treated with increasing concentrations of spermidine. (**C**) Relative viability analysis of cells from (B). (**D**) AlphaFold 3 prediction of the ATP13A3-AZ1 complex structures in human, fly, worm, and yeast. (**E**) Table of ipTM scores for all predicted complexes shown in (D). (**F**) Proposed model for ATP13A3-AZ1 mediated regulation of polyamine transport. Left: Under low polyamine concentrations ATP13A3 is directed to the plasma membrane. Increasing polyamine levels lead to expression of antizyme that binds and internalizes the transporter. Right: Disruption of this interaction causes polyamine toxicity through uncontrolled polyamine influx.

### ATP13A3-AZ binding is conserved from yeast to human

Since the regulation of ODC by AZ is well conserved among eukaryotes, we investigated whether ATP13A3 regulation was similarly conserved. We generated confident AlphaFold 3 models of ATP13A3-AZ complexes from fly, worm, and yeast (Fig. 5, D and E). The WHK and RF motifs of ATP13A3 and AZ1, respectively, were strictly conserved among vertebrates, but divergent in fly, worm, and yeast (fig. S5, D and E). This is consistent with the large ATP13A3-AZ1 interface which relies less on individual conserved residues and more on the overall fold. Notably, worms have two ATP13A3 orthologs, Catp5 and Catp6, both of which confidently bind AZ *in silico*. We note that yeast YPK9 is annotated as a vacuolar transporter, consistent with the prevailing view that ATP13A family members are predominantly vesicular. Whether the ATP13A3 orthologs also localize to the plasma membrane under polyamine-deficient conditions and how they maintain polyamine homeostasis in concert with other plasma membrane polyamine permeases in these organisms remains to be determined.

## Discussion

We identified ATP13A3 as a plasma membrane polyamine transporter whose localization is regulated by intracellular polyamine levels. Different cell lines showed varying localization properties under basal conditions, likely because intrinsic differences in polyamine biosynthesis lead to different polyamine uptake requirements. In all cell lines, however, DFMO treatment led to nearly exclusive plasma membrane localization. Mechanistically, ATP13A3 internalization requires AZ expression and binding to the transporter. Disruption of this interaction by either AZ silencing or mutating the binding interface induces cellular polyamine toxicity (Fig. 5F). These findings challenge the current model in which ATP13A3 acts primarily as a transporter in endolysosomal compartments, transporting endocytosed polyamines from the lumen to the cytoplasm.

Regulation of polyamine uptake by AZ was demonstrated more than 30 years ago across various cell lines. However, the molecular mechanism was never defined (*11, 12, 26*). Recently, several polyamine transporters were discovered in humans, including members of the ATP13A and SLC family such as the neuronal plasma membrane transporter SLC45A4 (*16, 17, 32*). However, the identity of the main plasma membrane polyamine transporter in metazoans and its regulation remained unknown (*14*). Our findings fill these gaps and provide a mechanistic explanation for how polyamine homeostasis is maintained through an interplay of biosynthesis and uptake. Evolution of the polyamine-sensing frameshift mechanism that governs AZ expression was likely a key step in regulating cellular polyamine levels. AZ binding to ODC regulates the rate-limiting step in polyamine biosynthesis, while its binding to ATP13A3 controls polyamine influx, coupling biosynthesis and transport in a single feedback circuit. The high evolutionary conservation of the ATP13A3-AZ complex and broad ATP13A3 expression levels suggest that ATP13A3 is the principal plasma membrane polyamine transporter - a hypothesis that requires validation across tissues and organisms.

Our findings also have therapeutic implications for diseases such as cancer or conditions such as aging. Elevated polyamine levels are a hallmark of cancer, whereas declining polyamine levels have been found in aging (*5, 33*). In both cases, cellular polyamine homeostasis is altered providing an opportunity for intervention in either direction. The ODC inhibitor DFMO is already used in high-risk neuroblastoma, but a key limitation of inhibiting polyamine synthesis is often observed compensatory upregulation of polyamine uptake. One polyamine depletion strategy therefore combines DFMO treatment with the PTI AMXT-1501, a combination currently being investigated in clinical trials for solid tumors (ClinicalTrials.gov: NCT07287917, NCT06465199). By identifying ATP13A3 as the regulated plasma membrane polyamine transporter, our work provides a means to target polyamine uptake. An alternative therapeutic strategy could involve disrupting the ATP13A3-AZ interaction to trap ATP13A3 at the plasma membrane. Appropriately fine-tuned, this could stabilize polyamine levels, for instance in aging, or drive them to cytotoxic concentrations for the treatment of cancer. ATP13A3 and its AZ-dependent regulation thus emerge as promising candidates for targeting polyamine homeostasis.

## Acknowledgments

We thank A. Carracedo for discussions on polyamine metabolism; K. Fröhlich for small-molecule LC-MS analysis; R. Girish for help in image quantification. Claude (Anthropic) was used for the analysis of clonogenic growth assays and for coding of Fiji and Rosetta scripts.

## Funding

Postdoctoral Fellowship of the Alexander-von-Humboldt Foundation (CM)

Postdoctoral Fellowship of the Engelhorn Foundation (CM)

EMBO Postdoctoral Fellowship (GCS)

SNF project grant 320030-227610 (MNH)

Balzan prize 2024 (MNH)

## Author contributions

Conceptualization: CM, MNH, DM Methodology: CM, GCS, MO, JKC, DR Investigation: CM, GCS, ABH, MO, JKC, DR Visualization: CM, GCS, MC

Funding acquisition: MNH, CM, GCS Supervision: MNH, CM

Writing – original draft: CM

Writing – review & editing: CM, GCS, DM, MNH

## Data, code, and materials availability

Proteomic data have been deposited to the ProteomeXchange Consortium (https://www.proteomexchange.org/) via the MassIVE partner repository (https://massive.ucsd.edu/) with MassIVE data set identifier MSV000102247 and ProteomeXchange identifier PXD080051. All other data are available in the main text or the supplementary materials.

## Supplementary Materials

Materials and Methods

Figs. S1 to S5

Table S1

## Materials and Methods

### Cell culture

Human liver cancer cell lines SNU-449, Hep3B, HuH-6, and HepG2 were gifted by Prof. Diego Calvisi (University of Greifswald, Germany), HuH-1 was gifted by Prof. Gerhard Christofori (University of Basel, Switzerland). HEK293T cells were obtained from ATCC. All cells were cultured in high glucose-containing DMEM (Sigma, #D5671) supplemented with 10% FBS, 2 mM glutamine, 0.1 mM non-essential amino acids (Gibco, #11140-035), 1x penicillin/streptomycin (Sigma, #P4333) and 250 µM aminoguanidine (Sigma, #396494), hereafter referred to as DMEM complete. Cells were incubated at 37 °C with 5% CO2 and regularly tested for mycoplasma contamination.

### Generation of stable cell lines

For stable expression of GFP-ATP13A2 and GFP-ATP13A3, both genes were cloned into the lentiviral vector pLenti-CMV-N-GFP DEST (Addgene plasmid #19732) using Gateway cloning. For doxycycline-inducible expression (iODC, iAZ1, iGFP) lentiviral pLV[Exp]-Puro-TRE>3xFLAG/gene_of_interest vectors were obtained from Vector Builder. For doxycycline-inducible knockdown of *OAZ1* pLV[shRNA]-TetR:IRES:Neo-U6/2xTetO>hOAZ1 (#1: ATGTTGTAATCGTGCAAATAACTCGAGTTATTTGCACGATTACAACAT, #2:ACAATCTTTCAGCTAACTTATCTCGAGATAAGTTAGCTGAAAGATTGT) plasmids were obtained from Vector Builder. For the expression of the Tet regulatory protein rtTA, required for achieving Tet-inducible expression, pLV[Exp]-Bsd-CMV>HA/rtTA was obtained from Vector Builder.

HEK293T cells were co-transfected with a lentiviral vector containing the gene of interest, psPAX2 (a gift from Didier Trono: Addgene plasmid #12260) and pCMV-VSV-G (a gift from Robert Weinberg: Addgene plasmid #8454). Supernatants containing lentiviral particles were collected 48 hours after transfection, pooled, and filtered (0.2 µm). Filtered supernatants were used to infect target cells. Infected cells were selected with 2 µg/ml puromycin (Gibco, #A11138-03) (GFP-A2 and GFP-A3), 20 µg/ml blasticidin (InvivoGen, #ant-bl-05) (rtTA), 1 mg/ml G418 (BioConcept, # 4-15F01-H) (iODC, iGFP, iAZ), or a combination. Selected cells were FACS-sorted for GFP to single cells or single-seeded by limited dilution into 96-well plates. Cells were propagated and tested for expression or knockdown by immunoblotting or real-time quantitative polymerase chain reaction (qPCR), respectively.

For CRISPR/Cas9-mediated knockout (KO) in SNU-449 cells, plasmids containing Cas9, a GFP reporter and three guide RNAs per target were purchased from Santa Cruz Biotechnology (USA) for ATP13A2 (sc-404770) and ATP13A3 (sc-413173). Cells were seeded in DMEM complete and the next day transfected with the described plasmids using jetPRIME (Sartorius, # 201000003). The cells were single clone FACS-sorted into 96-well plates using the GFP reporter 48 h post transfection. Cells were propagated and tested for KO by PCR. The PCR-amplified fragments were sequenced and successful clones were validated by Immunoblotting.

### Protein expression and purification for SEC-MALS

Human ATP13A3 nucleotide-binding domain (NTD, residues 500–726), an ATP13A3 NTD loop-swap mutant (LS; residues 690–702 replaced with the corresponding ATP13A2 sequence), and human OAZ1 (residues 91–228) were cloned into a pET10 Strep-TEV expression vector by VectorBuilder (vector IDs: VB251030-1382prq, VB251030-1360vsm, and VB251030-1395cbd; vectorbuilder.com). ATP13A3-NTD and OAZ1 constructs were subsequently modified by site-directed mutagenesis to incorporate a second Strep-tag sequence followed by a TEV cleavage site (for primers see Table S1).

All recombinant proteins were expressed in *E. coli* BL21(DE3) cells. Overnight starter cultures were diluted into 1 L of Luria-Bertani (LB) broth, and protein expression was induced with 0.5 mM isopropyl β-d-1-thiogalactopyranoside (IPTG) at OD₆₀₀ = 0.6. Cultures were grown overnight at 18 °C, harvested by centrifugation, and resuspended in lysis buffer (150 mM NaCl, 20 mM Tris-HCl pH 7.5, cOmplete protease inhibitor cocktail, 1 mM PMSF). Cells were lysed by sonication, and lysates were clarified by centrifugation at 15,000 × *g* for 45 min at 4 °C. Clarified lysates were incubated with 1 ml of equilibrated Strep-Tactin TACS agarose resin (IBA Lifesciences) for 2 hr at 4 °C with rotation. The resin was washed with 30 ml of wash buffer (150 mM NaCl, 20 mM Tris-HCl pH 7.5), and bound proteins were eluted by 10 min incubation in wash buffer supplemented with 1 mM d-desthiobiotin (IBA Lifesciences). Eluted proteins were further purified by size-exclusion chromatography (SEC) on a Superdex 200 10/300 GL column (Cytiva) equilibrated in wash buffer. All recombinant proteins eluted as a monodisperse peak.

### Site-directed mutagenesis

Site-directed mutagenesis was carried out using the polymerase chain reaction (*34*) or the Q5 Site-Directed Mutagenesis Kit (NEB, E0554S) following the manufacturer’s protocol. All constructs were verified by whole-plasmid sequencing (Microsynth). Primers are listed in Table S1.

### Clonogenic growth assays and crystal violet staining

500 cells were seeded into 12-well plates in DMEM complete medium. Polyamine synthesis or uptake inhibitors, polyamines were added directly to the medium on the day of seeding, while DMSO (0.1%) or Doxycycline (1 µg/ml) were added the next day. Cells were incubated for 7 days before staining. To visualize colony formation, cells were stained with crystal violet (2% (v/v) crystal violet in 20% (v/v) methanol).

### Immunoblots

Cells from human liver cancer cell lines were lysed in M-PER (ThermoFisher Scientific, #78501) supplemented with cOmplete inhibitor cocktail (Roche, #11836170001) and PhosSTOP (Roche, #04906837001). Protein concentration was determined by Pierce BCA assay (ThermoFisher Scientific, #23225), and equal amounts of protein were separated by SDS-PAGE, and transferred onto nitrocellulose membranes (GE Healthcare). Membranes were blocked in 5 % milk in Tris buffered saline with 0.1% Tween20 (TBST) and incubated with primary antibody overnight. Secondary antibody was added for 1 h and blots were incubated with SuperSignal Western Blot Substrate Pico (ThermoFisher Scientific, #34580) or Femto (ThermoFisher Scientific, #34095) and imaged using a Fusion FX imaging system (Vilber). Antibodies used in this study were as follows: Actin (Millipore, #MAB1501), FLAG (Sigma, #F1804), GPF (Roche, #11814460001), ATP13A2 (Sigma-Aldrich, #A3361), ATP13A3 (Sigma-Aldrich, #HPA029471 (fig. S1); (Novus Biologicals/ Bio-techne, #NBP3-06070) (all other figures)), AZ1 (Boster, #A06486-4), mouse anti-rabbit (Jackson ImmunoResearch, #211-032-171), goat anti-mouse (Jackson ImmunoResearch, #115-035-174).

### RNA isolation and quantitative real-time PCR

Total RNA was isolated using the RNeasy Kit (QIAGEN). RNA was reverse transcribed using iScript cDNA Synthesis Kit (Bio-Rad). Quantitative real-time PCR (qPCR) analysis was performed using Fast SYBR Green (Applied Biosystems) and a q3TOWER (AnalytikJena). Relative expression levels were determined using the ΔΔCt method. For each gene at least three independent biological replicates were used. The primer pairs are shown in Table S1.

### siRNA-mediated knockdown

For siRNA-mediated knockdown, cells were seeded in DMEM complete medium and transfected the next day with 50 nM siRNA using DharmaFECT4 (Horizon Discovery, UK), in OptiMEM. Medium was exchanged the following day, and cells were incubated for 48 h prior to harvest and analysis by qPCR. The following siRNAs were used in this study: *SLC3A2* (pooled) #L-003542-00-0005; *ATP13A3* (pooled) #L-008131-01-0005; *ATP13A2* (single siRNA) #J-008601-08-0005; non-targeting #D-001810-01-05 (all Horizon Discovery); *OAZ1* (pooled) # sc-105112 (Santa Cruz Biotechnology).

### 3H-putrescine and ^3^H-spermidine uptake

Cells were seeded in DMEM complete and supplemented, as indicated, with 1 mM DFMO for 48 hours to induce polyamine depletion. Polyamine transport inhibitor (PTI) AMXT-1501 was added at 1 µM concentration for 1 h before the start of the assay. Cells were pre-treated for 2 h with 10 µM polyamines, 10 mM arginine or 1 mM ornithine, as indicated. Putrescine [2,3-^3^H(N)] dihydrochloride or Spermidine [terminal methylene-^3^H] was diluted in DMEM complete, supplemented with cold putrescine or spermidine, respectively and added to the cells at a final concentration of 1 µM polyamine (100 nCi/ml for putrescine, 100 nCi/ml for spermidine) for 15 min at 37°C. Cells were placed on ice, washed twice with cold PBS and lysed with 1 M HCl. The lysate was directly transferred to scintillation counter tubes containing Ultima Gold liquid scintillation cocktail (Revivity #6013321) and measured using a scintillation counter (PerkinElmer).

### Sample preparation of GFP-A3 pull-down

SNU-449 A3^-/-^ GFP-A3 cells were seeded at 1×10⁴ cells/mL in 6 cm dishes (4 mL per dish, 8 dishes per experiment). Four dishes were treated with putrescine (10 µM, 6 h) and four served as untreated controls. Cells were washed with PBS and lysed in 500 µL lysis buffer containing detergent per dish. Plates were frozen at −80°C and subsequently scraped in lysis buffer and transferred to 2 mL tubes. Lysates were sonicated with 5 short pulses (1 s each) in the cold room and incubated for 1 h at 4°C with rotation. Lysates were cleared by centrifugation at 18,000 g for 30 min at 4°C. GFP-Trap beads (ChromoTek, #gta) were washed three times with lysis buffer and re-suspended in lysis buffer. Cleared lysates were added to the beads and incubated overnight at 4°C with rotation. For PTM profiling, bead-bound material was transferred to columns, washed three times with wash buffer (20 s, 50 g), and eluted by boiling in elution buffer (5% SDS, 10 mM TCEP, 100 mM TEAB) at 95°C for 10 min. Eluates were processed using the S-trap protocol with 1.5 h trypsin digestion. Peptides were dried by vacuum centrifugation and stored at −20°C until mass spectrometric analysis. For interactome analysis, bound proteins were eluted by boiling in SDS sample buffer and analyzed by mass spectrometry. All samples were further processed and analyzed by mass spectrometry as described below. Lysis buffer consisted of 20 mM Tris/HCl pH 7.5, 300 mM NaCl, 1 mM EDTA, 1 mM DTT, 10% glycerol, and cOmplete protease inhibitor cocktail (Roche), supplemented with 1% DDM and 0.1% CHS as detergents. Wash buffer was prepared identically but with reduced detergent concentrations of 0.03% DDM and 0.003% CHS.

### LC-MS analysis of GFP-A3 pull-down

Proteins were eluted by incubation for 10 min at 95°C in lysis buffer (5% SDS, 10mM TCEP, 0.1 M TEAB), alkylated in 20 mM iodoacetamide for 30 min at 25°C and digested using S-Trap™ micro spin columns (Protifi) according to the manufacturer’s instructions. Shortly, 12 % phosphoric acid was added to each sample (final concentration of phosphoric acid 1.2%) followed by the addition of S-trap buffer (90% methanol, 100 mM TEAB pH 7.1) at a ratio of 6:1. Samples were mixed by vertexing and loaded onto S-trap columns by centrifugation at 4000 g for 1 min followed by three washes with S-trap buffer. Digestion buffer (50 mM TEAB pH 8.0) containing sequencing-grade modified trypsin (1/25, w/w; Promega, Madison, Wis-consin) was added to the S-trap column and incubated for 1h at 47°C. Peptides were eluted by the consecutive addition and collection by centrifugation at 4000 g for 1 min of 40 ul digestion buffer, 40 uL of 0.2% formic acid and finally 35 uL 50% acetonitrile, 0.2% formic acid. Samples were dried under vacuum and stored at - 20°C until further use. Dried peptides were resuspended in 0.1% aqueous formic acid and subjected to LC–MS/MS analysis using a Orbitrap Fusion Lumos Mass Spectrometer fitted with an EASY-nLC 1200 (both Thermo Fisher Scientific) and a custom-made column heater set to 60°C. Peptides were resolved using a RP-HPLC column (75μm × 30cm) packed in-house with C18 resin (ReproSil Saphir 100 C18, 1.5 um resin; Dr. Maisch GmbH) at a flow rate of 0.2 ul/min. The following gradient was used for peptide separation: from 2% B to 8% B over 5 min to 25% B over 40 min to 35% B over 15 min to 95% B over 2 min followed by 18 min at 95% B. Buffer A was 0.1% formic acid in water and buffer B was 80% acetonitrile, 0.1% formic acid in water. The mass spectrometer was operated in DDA mode with a cycle time of 3 seconds between master scans. Each master scan was acquired in the Orbitrap at a resolution of 240,000 FWHM (at 200 m/z) and a scan range from 375 to 1600 m/z followed by MS2 scans of the most intense precursors in the Orbitrap at a resolution of 30,000 FWHM (at 200 m/z) with isolation width of the quadrupole set to 1.4 m/z. Maximum ion injection time was set to 50ms (MS1) and 54 ms (MS2) with an AGC target set to 1e6 and 5e4, respectively. Only peptides with charge state 2 – 5 were included in the analysis. Monoisotopic precursor selection (MIPS) was set to Peptide, and the Intensity Threshold was set to 2.5e4. Peptides were fragmented by HCD (Higher-energy collisional dissociation) with collision energy set to 35%, and one microscan was acquired for each spectrum. The dynamic exclusion duration was set to 30s. The acquired raw files were searched using both MSFragger and Metamorpheus. MSFragger (v. 4.3) implemented in FragPipe (v. 23.1) was used to search raw files against a H. sapiens database (consisting of 20360 protein sequences downloaded from Uniprot on 20220222) and 392 commonly observed contaminants using the “LFQ-MBR” workflow with the following changes: i) carbamidomethylation of Cysteine was not used as a fixed modification, ii) ionquant normalization was switched off. Metamorpheus was used for PTM discovery against the same H sapiens database described above using default settings.

### TMT labelling and LC-MS analysis of SNU-449 A3^-/-^ cells

Washed cell pellets were resuspended in lysis buffer (5% SDS, 10mM TCEP, 0.1 M TEAB) and lysed by sonication using a PIXUL Multi-Sample Sonicator (Active Motif) with Pulse set to 50, PRF to 1, Process Time to 10 min and Burst Rate to 20 Hz. Lysates were incubated for 10 min at 95°C, alkylated in 20 mM iodoacetamide for 30 min at 25°C and proteins digested using S-Trap™ micro spin columns (Protifi) according to the manufacturer’s instructions. Shortly, 12 % phosphoric acid was added to each sample (final concentration of phosphoric acid 1.2%) followed by the addition of S-trap buffer (90% methanol, 100 mM TEAB pH 7.1) at a ratio of 6:1. Samples were mixed by vertexing and loaded onto S-trap columns by centrifugation at 4000 g for 1 min followed by three washes with S-trap buffer. Digestion buffer (50 mM TEAB pH 8.0) containing sequencing-grade modified trypsin (1/25, w/w; Promega, Madison, Wisconsin) was added to the S-trap column and incubated for 1h at 47 °C. Peptides were eluted by the consecutive addition and collection by centrifugation at 4000 g for 1 min of 40 ul digestion buffer, 40 ul of 0.2% formic acid and finally 35 ul 50% acetonitrile, 0.2% formic acid. Samples were dried under vacuum and stored at-20 °C until further use.

Sample aliquots comprising 10 μg of peptides were labeled with isobaric tandem mass tags (TMTpro 16-plex, Thermo Fisher Scientific). Peptides were resuspended in 10 μl labeling buffer (2 M urea, 0.2 M HEPES, pH 8.3) by sonication and 2.5 μl of each TMT reagent were added to the individual peptide samples followed by a 1 h incubation at 25°C shaking at 500 rpm. To quench the labelling reaction, 0.75 μl aqueous 1.5 M hydroxylamine solution was added and samples were incubated for 5 min at 25°C shaking at 500 rpm followed by pooling of all samples. The pH of the sample pool was increased to 11.9 by adding 1 M phosphate buffer (pH 12) and incubated for 20 min at 25°C and 500 rpm shaking to remove TMT labels linked to peptide hydroxyl groups. Subsequently, the reaction was stopped by adding 2 M hydrochloric acid until a pH < 2 was reached. Finally, peptide samples were further acidified using 5 % TFA, desalted using BioPureSPN MACRO™ SPE cartridges (Nest group) according to the manufacturer’s instructions and dried under vacuum.

TMT-labeled peptides were fractionated by high-pH reversed phase separation using a XBridge Peptide BEH C18 column (3.5 µm, 130 Å, 1 mm x 150 mm, Waters) on an Ultimate 3000 system (Thermo Scientific). Peptides were loaded on column in buffer A and the system was run at a flow of 42 µl/min. The following gradient was used for peptide separation: from 2% B to 15% B over 3 min to 45% B over 59 min to 80% B over 3 min followed by 9 min at 80% B then back to 2% B over 1 min followed by 15 min at 2% B. Buffer A was 20 mM ammonium formate in water, pH 10 and buffer B was 20 mM ammonium formate in 90% acetonitrile, pH 10. Elution of peptides was monitored with a UV detector (205 nm, 214 nm) and a total of 36 fractions were collected, pooled into 12 fractions using a post-concatenation strategy as previously described (Wang et al., 2011) and dried under vacuum.

Dried peptides were resuspended in 0.1% aqueous formic acid and subjected to LC–MS/MS analysis using an Orbitrap Eclipse Tribrid Mass Spectrometer fitted with a Ultimate 3000 nano system and a FAIMS Pro interface (all Thermo Fisher Scientific) and a custom-made column heater set to 60°C. Peptides were resolved using a RP-HPLC column (75 μm × 30 cm) packed in-house with C18 resin (ReproSil-Pur C18–AQ, 1.9 μm resin; Dr. Maisch GmbH) at a flow rate of 0.3 μl/min. The following gradient was used for peptide separation: from 2% B to 12% B over 5 min to 30% B over 70 min to 50% B over 15 min to 95% B over 2 min followed by 18 min at 95% B then back to 2% B over 2 min followed by 18 min at 2% B. Buffer A was 0.1% formic acid in water and buffer B was 80% acetonitrile, 0.1% formic acid in water.

The mass spectrometer was operated in DDA mode with a cycle time of 3 s between master scans. Throughout each acquisition, the FAIMS Pro interface switched between CVs of −40 V and −70 V with cycle times of 1.5 s and 1.5 s, respectively. MS1 spectra were acquired in the Orbitrap at a resolution of 120,000 and a scan range of 400 to 1600 m/z, AGC target set to “Standard” and maximum injection time set to “Auto”. Precursors were filtered with precursor selection range set to 400–1600 m/z, monoisotopic peak determination set to “Peptide”, charge state set to 2 to 6, a dynamic exclusion of 45 s, a precursor fit of 50% in a window of 0.7 m/z and an intensity threshold of 5e3.

Precursors selected for MS2 analysis were isolated in the quadrupole with a 0.7 m/z window and collected for a maximum injection time of 35 ms with AGC target set to “Standard”. Fragmentation was performed with a CID collision energy of 30% and MS2 spectra were acquired in the IT at scan rate “Turbo”.

MS2 spectra were subjected to RTS using a human database containing 20362 entries downloaded from Uniprot on 20200417 using the following settings: enzyme was set to “Trypsin”, TMTpro16plex (K and N-term) and Carbamidomethyl (C) were set as fixed modification, Oxidation (M) was set as variable modifications, maximum missed cleavages was set to 1 and maximum variable modifications to 2. Maximum search time was set to 100 ms, the scoring threshold was set to 1.4 XCorr, 0.1 dCn, 10 ppm precursor tolerance, charge state 2 and “TMT SPS MS3 Mode” was enabled. Subsequently, spectra were filtered with a precursor selection range filter of 400–1600 m/z, precursor ion exclusion set to 25 ppm low and 25 ppm high and isobaric tag loss exclusion set to “TMTpro”. MS/MS product ions of precursors identified via RTS were isolated for an MS3 scan using the quadrupole with a 2 m/z window and ions were collected for a maximum injection time of 200 ms with a normalized AGC target set to 200%. SPS was activated and the number of SPS precursors was set to 10. Isolated fragments were fragmented with normalized HCD collision energy set to 55% and MS3 spectra were acquired in the orbitrap with a resolution of 50,000 and a scan range of 100 to 500 m/z.

The acquired raw-files were analysed using the SpectroMine software (Biognosis AG, Schlieren, Switzerland). Spectra were searched against a human database consisting of 20372 protein sequences (downloaded from Uniprot on 20220222). Standard Pulsar search settings for TMTpro18 (“TMTpro18_Quantification”) were used and resulting identifications and corresponding quantitative values were exported on the PSM level using the “Export Report” function. Acquired reporter ion intensities were employed for automated quantification and statistical analysis using the in-house developed SafeQuant R script (v2.3) (Ahrné et al., 2016). This analysis included adjustment of reporter ion intensities, global data normalization by equalizing the total reporter ion intensity across all channels, data imputation using the knn algorithm, summation of reporter ion intensities per protein and channel and calculation of protein abundance ratios. To meet additional assumptions (normality and homoscedasticity) underlying the use of linear regression models and t-tests, MS-intensity signals were transformed from the linear to the log-scale. The summarized protein expression values were used for statistical testing of between condition differentially abundant proteins. Here, empirical Bayes moderated t-tests were applied, as implemented in the R/Bioconductor limma package (http://bioconductor.org/packages/release/bioc/html/limma.html). The resulting per protein and condition comparison p-values were adjusted for multiple testing using the Benjamini-Hochberg method.

### Targeted LC-MS analysis of SNU-449 A2^-/-^ cells

The following 11 peptide sequences were selected to target ATP13A2 (Q9NQ11, AT132_HUMAN): TGTLTEDGLDVMGVVPLK2, VVALASKPLPTVPSLEAAQQLTR3, DTAQLHK2, HLALSGPTFGIIVK3, ELAEQPWPPLPAGPLR2, AIYGPNVISIPVK2, YYLFQGQR2, VPLNEIVIR2, VLVQGTVFAR2, LQDTPVGDPMDLK2, TALPEGLGPYC[+57.021464]AETHRR4 (charge state indicated as number at the end of the respective sequence). A mass isolation list containing the masses of these peptides at the indicated charge states was exported from Skyline and imported into the Orbitrap Fusion Lumos operating software for targeted analysis.

Peptide samples were resuspended in 0.1% aqueous formic acid and subjected to LC–MS/MS analysis using a Orbitrap Fusion Lumos Mass Spectrometer fitted with an EASY-nLC 1200 (both Thermo Fisher Scientific) and a custom-made column heater set to 60°C. Peptides were resolved using a RP-HPLC column (75 μm × 37 cm) packed in-house with C18 resin (ReproSil-Pur C18–AQ, 1.9 μm resin; Dr. Maisch GmbH) at a flow rate of 0.2 ul/min. A linear gradient ranging from 5% buffer B to 45% buffer B over 60 minutes was used for peptide separation. Buffer A was 0.1% formic acid in water and buffer B was 80% acetonitrile, 0.1% formic acid in water. The mass spectrometer was operated in PRM mode with a maximum cycle time of 2.7 seconds between master scans. Each master scan was acquired in the Orbitrap at a resolution of 120,000 FWHM (at 200 m/z), scan range from 300 to 1600 m/z, maximum injection time of 100 ms and normalized AGC target set to 250%. MS2 scans were acquired in the Orbitrap in centroid mode at a resolution of 120,000 FWHM (at 200 m/z), a scan range from 150 to 1500 m/z, maximum injection time of 246 ms, normalized AGC target set to 200% with an ion isolation window of 0.4 m/z. Peptides were fragmented by HCD (Higher-energy collisional dissociation) with collision energy set to 35%, loop control was set to “All” and one microscan was acquired for each spectrum. The acquired raw files were searched using DIA-NN (v.1.8.1) in “library-free search” mode against a Homo sapiens database (consisting of 20360 protein sequences downloaded from Uniprot on 20220222) and 393 commonly observed contaminants using default library-free search settings with the following modifications: 1) oxidation of methionine was set as variable modification, 2) Protein inference was set to “Protein names (from FASTA)”.

### AlphaFold 3 modeling

AlphaFold 3 was used to predict interactions between different ATP13A and antizyme proteins across different organisms (*28*). Protein sequences were used as input, and the analyses were directly run on the AlphaFold server through the web-interface. Coordinate files were used for further analysis and in structure visualization. Reported interface predicted template modeling (ipTM) values were used to assess prediction confidence. Values <0.6 suggest likely failed predictions, 0.6-0.8 are a gray zone where predictions could be correct, while values >0.8 represent confident predictions (*28*).

### Targeted metabolomics of SNU-449 cells

Cells for targeted metabolomics of polyamines were depleted of endogenous polyamines by treating the cells with 1 mM DFMO for 48 h. The cells were then supplemented with 100 µM of individual polyamine for 1 h before washed with PBS. The metabolites were extracted by adding 1 ml of 80% ethanol to the dish and the suspension was frozen at-80 °C for at least 30 min. Extracts were separated from cell debris by centrifuging at 20’00 g at 4°C for 10 minutes, dried in a SpeedDry Vacuum Concentrator (Christ). Dried samples were reconstituted in 100 μL of 40% acetonitrile, 0.1% formic acid in MilliQ water. A dilution of 1:10 was performed using 10% acetonitrile, 0.1% formic acid in MilliQ water. Samples were analyzed using a Vanquish HPLC (Thermo Fisher) coupled to a Compact II (Bruker). 5µl of sample were injected and separated using an ACQUITY Premier BEH C18 1.7µm 2.1 x 100mm column (Waters) heated to 40°C and employing a 5min gradient ranging from 2 to 100% buffer B at 200µl/min flow. Buffer A consisted of 0.1% formic acid. Buffer B consisted of 80% acetonitrile, 0.1% formic acid. The mass spectrometer was operated in positive mode, using 500V end plate offset, 3500V capillary voltage, 0.4 bar Nebulizer pressure, 4 l/min dry gas and 180°C dry temperature as source settings. The tune parameters were set to 0 eV isCIDenergy, 70 V hexapole voltage, 2 eV quadrupole Ion energy, 100 m/z low mass and a pre pulse storage of 2µs was employed. Only MS1 spectra were recorded at 1 Hz. Data Analysis was performed using Skyline Version 21.1 in small molecules mode. The mass tolerance was set to 50 ppm.

### Sample preparation for fluorescence microscopy

Cells were plated in equal numbers on cover slips (VWR, #631-0147, No. 1.5) and cultured for 48 h in complete growth medium before treatment. Where indicated, cells received the relevant treatment (2 h 10 µM putrescine, 2 h 10 mM L-arginine, 2 h 1 mM L-ornithine, or 48 h 1 mM DFMO with or without a 2 h 10 µM polyamine pulse on the day of harvest or 30 h DMSO/doxycycline induction as described above) before being processed for imaging. Immediately before fixation, plasma membranes were labelled live with CellBrite Steady 640 (Biotium, #30107) at 1:1000 in PBS for 15 min at 37 °C. Cells were then fixed and counterstained in one step with 4 % paraformaldehyde (Electron Microscopy Sciences #15710) in PBS containing NucBlue Live ReadyProbes Reagent (Hoechst 33342; Thermo Fisher Scientific, #R37605) at 2 drops/mL for 10 min at 37 °C, followed by two PBS washes. For samples imaged without primary antibody staining (GFP fluorescence only), coverslips were mounted on glass slides with Vectashield Antifade Mounting Medium (Vector Laboratories, #H-1000) and sealed with nail varnish.

### Antibody staining for ICC experiments

For antibody-based detection of intracellular markers, fixed cells were permeabilized with 0.1 % Triton X-100 in PBS for 10 min at room temperature and blocked with 5 % bovine serum albumin (BSA) in PBS for 30 min at room temperature. Primary antibodies were diluted to 1:200 in 5 % BSA/PBS and applied overnight at 4 °C in a humidified chamber. Coverslips were washed three times for 5 min each with PBS and incubated with the appropriate Alexa Fluor 568-conjugated secondary antibody at 1:500 in 5 % BSA/PBS for 1 h at room temperature protected from light. After three additional 5 min PBS washes, coverslips were mounted and sealed as described above.

The following primary antibodies were used for immunofluorescence: mouse monoclonal anti-EEA1 (BD Biosciences # 610457; 1:200), rabbit monoclonal anti-GM130 (Cell Signaling, #1240S; 1:200), and rabbit monoclonal anti-LAMP1 (Cell Signaling Technology, clone D2D11 # 9091; 1:200). The secondary antibodies were goat anti-mouse IgG Alexa Fluor 568 (Invitrogen / Thermo Fisher Scientific, #A11004; 1:500), used for the mouse-host EEA1 and Rab11 primaries, and goat anti-rabbit IgG Alexa Fluor 568 (Invitrogen / Thermo Fisher Scientific, #A11036; 1:500), used for the rabbit-host GM130 and LAMP1 primaries.

### Image acquisition

Cells were imaged using a spinning disk Nikon Ti2 Crest X-Light V3 microscope in 60x oil-immersed objective (Plan Apo N). OMERO (Open Microscopy Environment) was used for data management and figure preparation.

### Image quantification for fluorescence microscopy

Image quantification was performed in Image J (Fiji) (*35*) using a custom script written in Jython. For each image, the middle z-slice was selected for analysis. Cell segmentation was performed using Cellpose with the cyto2 pretrained model (*36*), using the DAPI channel as a nuclear marker and the plasma membrane channel as the cytoplasmic guide channel. Segmentation parameters were: cell diameter 20 µM (HepG2) and 40 µM (all other cells), cell probability threshold 0, and flow threshold 0.4.

To quantify plasma membrane-associated protein enrichment, two compartment masks were derived from the Cellpose segmentation label image using morphological operations implemented in MorphoLibJ (*37*).A membrane ring mask was generated by subtracting a label-eroded and label-dilated image (erosion and dilation radii: 10 px for HepG2, 20 px for all other cells). Labels present in the membrane ring mask but absent after erosion, corresponding to cells too small to survive erosion, were removed to ensure one-to-one pairing between cell body and membrane ring ROIs. ROIs were converted from label images using the Labels2CompositeRois function of the BIOP Fiji plugin suite.

Mean GFP fluorescence intensity was measured independently within the cell body ROI and the corresponding membrane ring ROI for each segmented cell. The membrane-to-cell ratio was calculated as the mean intensity of the membrane ring divided by the mean intensity of the cell body, providing a measure of plasma membrane enrichment relative to total cellular fluorescence.

For colocalization analysis, pixel-wise metrics were computed between selected channel pairs on the middle z-slice. Pearson’s correlation coefficient was calculated across all pixels of the image to assess linear intensity correlation.

### Statistics

The investigators were not blinded to the treatment groups. Data are shown as mean ± SD or median (violin plots). Sample numbers are indicated in each figure legend. For cell culture experiments, n indicates the number of biological replicates. For quantification of fluorescence microscopy at least 10 images from three biological replicates were analyzed (>50 cells/condition). To determine the statistical significance unpaired Student’s t test, one-way-ANOVA, or two-way-ANOVA were performed using GraphPad Prism 11. A p value of less than 0.05 was considered statistically significant.

### Computational Modeling and Interface Energy Analysis

#### Initial structure preparation and relaxation

The predicted AlphaFold 3 model of the human ATP13A3–AZ1Δ1–90 complex was subjected to all-atom refinement using the FastRelax protocol implemented in the Rosetta molecular modeling suite (version 2019.21.60746). Refinement was performed using the ref2015_cart energy function with Cartesian minimization enabled. Twenty independent relaxation trajectories were generated to account for stochastic variation in side-chain packing and energy minimization. The resulting ensemble was evaluated using Rosetta energy metrics and interface quality descriptors calculated with InterfaceAnalyzer. A representative relaxed model was selected based on favorable interface binding energy (dG_separated), interface packing (packstat), and a low number of buried unsatisfied hydrogen bonds (delta_unsatHbonds). This refined structure was subsequently used as the common starting model for all computational mutagenesis analyses.

#### Identification of interface residues

Protein–protein interface residues were identified using the Rosetta InterfaceAnalyzer application. Residues contributing to the interaction surface were defined based on burial upon complex formation and changes in solvent-accessible surface area. The resulting interface residue set was used for subsequent computational mutagenesis analyses.

### Computational alanine scanning

To identify energetic hotspot residues within the protein–protein interface, mutant structures were generated from the selected wild-type model and refined using a local relaxation protocol. Structural flexibility was restricted to residues within a defined neighborhood surrounding the mutated position(s) using Rosetta MoveMaps. For multi-residue mutants, the relaxation region was defined as the union of local neighborhoods surrounding all mutated residues. Cartesian minimization with the ref2015_cart energy function was performed without global coordinate restraints. Twenty independent relaxation trajectories were generated for each construct prior to interface-energy calculations.

#### Combinatorial mutagenesis of ATP13A3LS

Selected combinations of mutations were introduced simultaneously into the protein sequence. Mutant complexes were subjected to the same relaxation and interface-analysis workflow as the wild-type structure. Twenty independently relaxed decoys were generated and analyzed for each multi-residue mutant construct.

### Interface Energy Calculations

Binding energetics were assessed using Rosetta InterfaceAnalyzer. The primary metric used to evaluate interaction strength was the separated binding energy (dG_separated), which estimates the energetic contribution of complex formation by comparing the energy of the bound complex to that of the separated binding partners.

Additional interface descriptors were recorded for quality assessment, including buried solvent-accessible surface area (dSASA_int), interface packing statistics (packstat), numbers of buried unsatisfied hydrogen bonds (delta_unsatHbonds), and intermolecular hydrogen-bond counts.

#### Calculation of Mutation Effects

The energetic effect of each mutation was quantified as the difference between mutant and wild-type interface binding energies:

ΔΔG = ⟨dG_separated(mutant)⟩ − ⟨dG_separated(wild-type)⟩, where ⟨dG_separated⟩ denotes the mean value obtained from the ensemble of independently generated decoys. Positive ΔΔG values indicate a predicted destabilization of the protein–protein interaction, whereas negative values indicate a predicted stabilization relative to the wild-type complex.

### Statistical Analysis

For each wild-type and mutant construct, 20 independent Rosetta trajectories were generated and analyzed. Reported interface energies are presented as mean ± standard deviation (SD) calculated across the complete ensemble of decoys.

### Conservation analysis

P5B ATPase protein family and OAZ1 sequences were retrieved from UniProt. All sequences were aligned using the Clustal Omega server (*38*) and visualized using ESPript 3.0 (*39*).

---

**Fig. S1.**
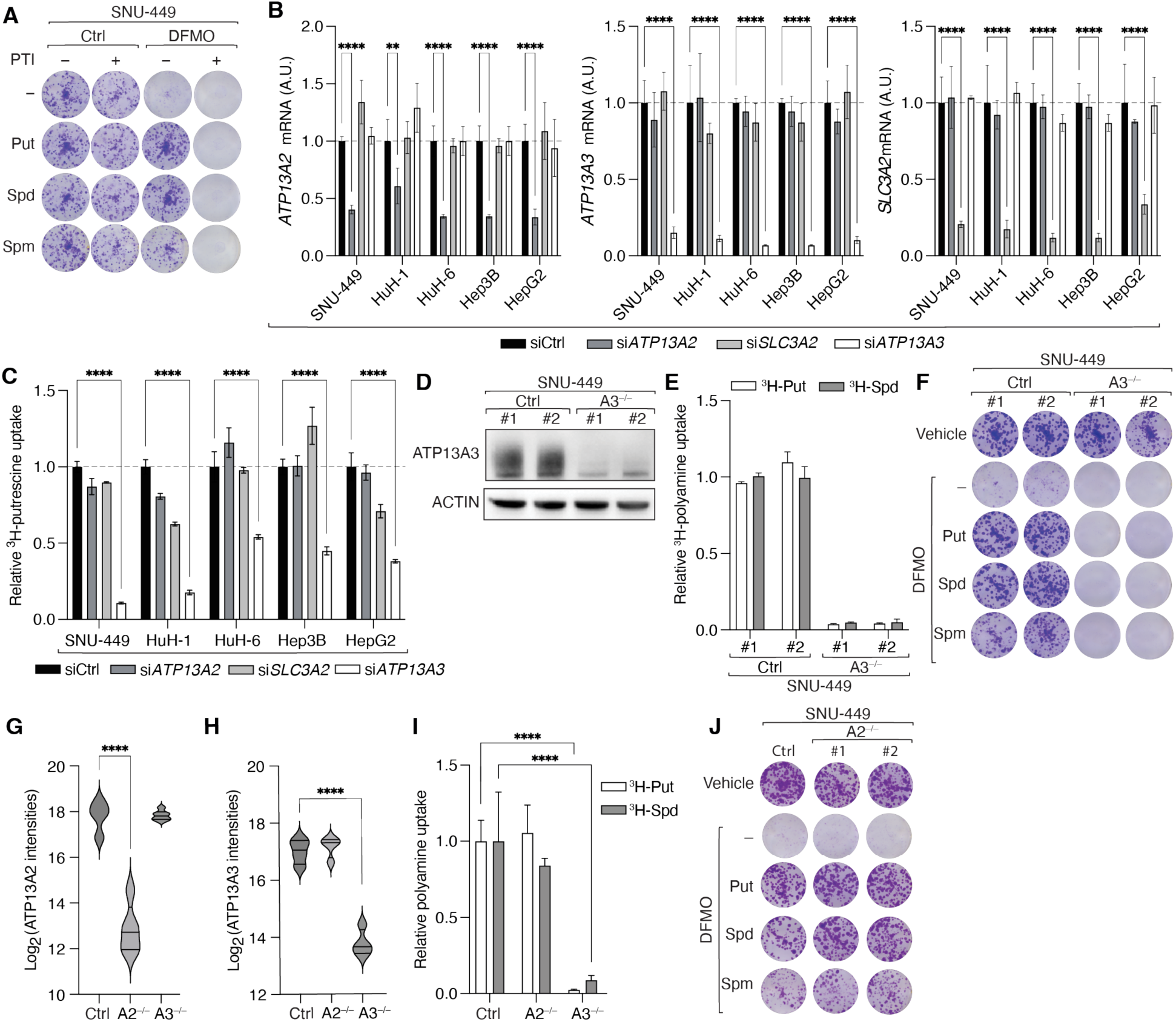
ATP13A3 is the main polyamine transporter in liver cells. **(A)** Representative clonogenic growth assay of SNU-449 cells treated with 1 mM DFMO, individual 1 µM polyamines, and 1 µM polyamine transporter inhibitor (PTI). **(B)** Relative gene expression of *ATP13A2*, *ATP13A3*, and *SLC3A2* in SNU-449, HuH-1, HuH-6, Hep3B, HepG2 cells upon treatment with siCtrl, si*ATP13A2*, si*ATP13A3*, or si*SLC3A2*. **(C)** Relative ^3^H-putrescine uptake into cells from (B) treated as in (B). n=4. **(D)** Immunoblots for ATP13A3 of two SNU-449 A3^-/-^ clones. ACTIN serves as loading control. **(E)** Relative ^3^H-polyamine uptake into cells from (D). n=4. **(F)** Representative clonogenic growth of cells from (D) treated as in (A). **(G** and **H)** Validation of SNU-449 A2^-/-^ and A3^-/-^ knockout clones using MS. n=4 clones. **(I)** Relative ^3^H-polyamine uptake into SNU-449 A2^-/-^ or A3^-/-^ cells. n=4 clones **(J)** Representative clonogenic growth assay of two SNU-449 A2^-/-^clones treated as in (A). ****p < 0.0001 by one-way-ANOVA (G, H) or two-way-ANOVA (B, C, E, I).

**Fig. S2.**
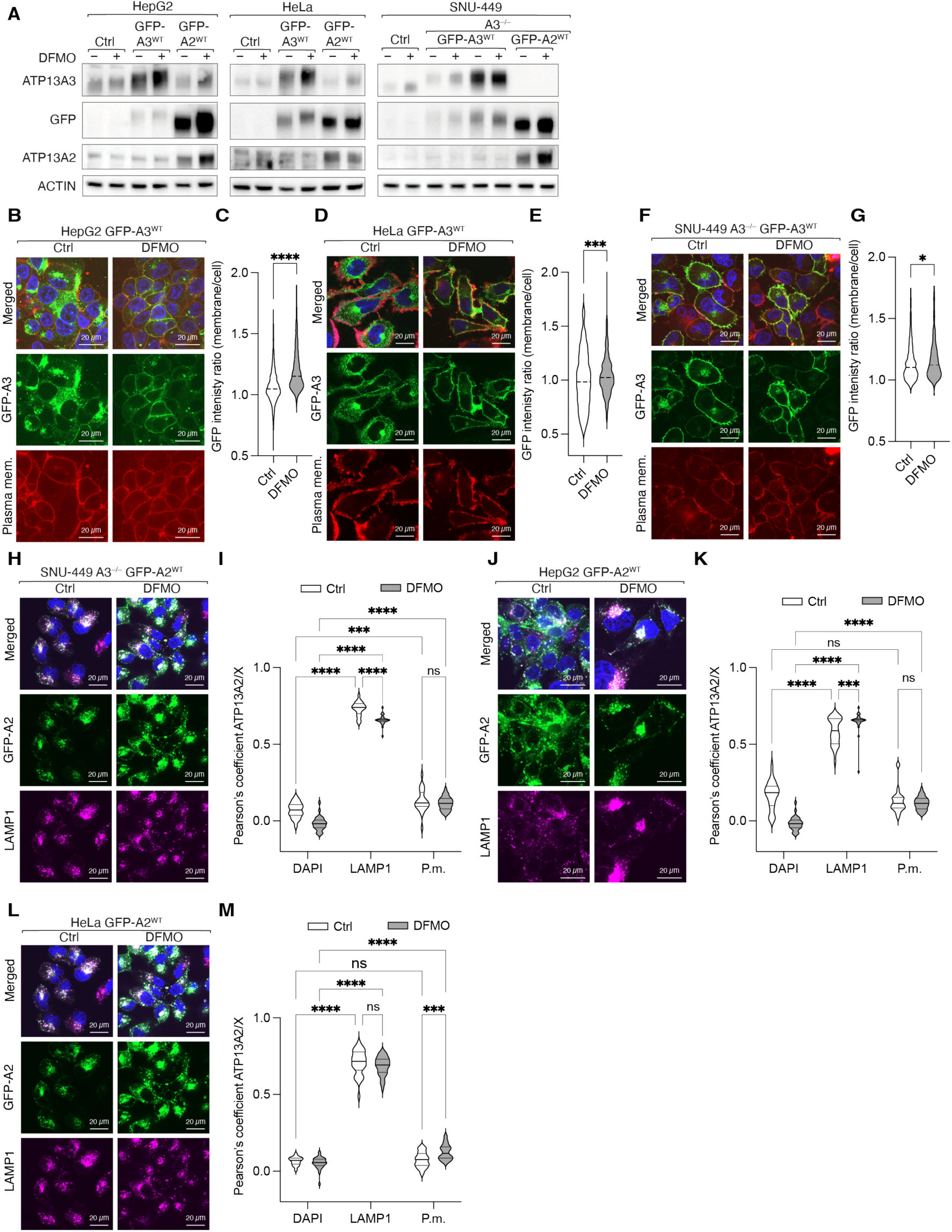
ATP13A3 but not ATP13A2 changes cellular localization upon changes in polyamine levels. (**A**) Immunoblots of ATP13A3, GFP, and ATP13A2 in HepG2 control, GFP-A3^WT^, and GFP-A2^WT^ cells; HeLa, GFP-A3^WT^, and GFP-A2^WT^; and SNU-449 control, A3^-/-^ GFP-A3^WT^ and A3^-/-^ GFP-A2^WT^ cells. ACTIN serves as loading control. (**B** to **M**) Fluorescence microscopy of GFP-A3^WT^ or GFP-A2^WT^ in cells from (A) in control or polyamine-depleted conditions (1 mM DFMO for 2 days). *p < 0.05, **p < 0.01, ***p < 0.001, ****p < 0.0001 by t-test (C, E, G) and two-way-ANOVA (I, K, M).

**Fig. S3.**
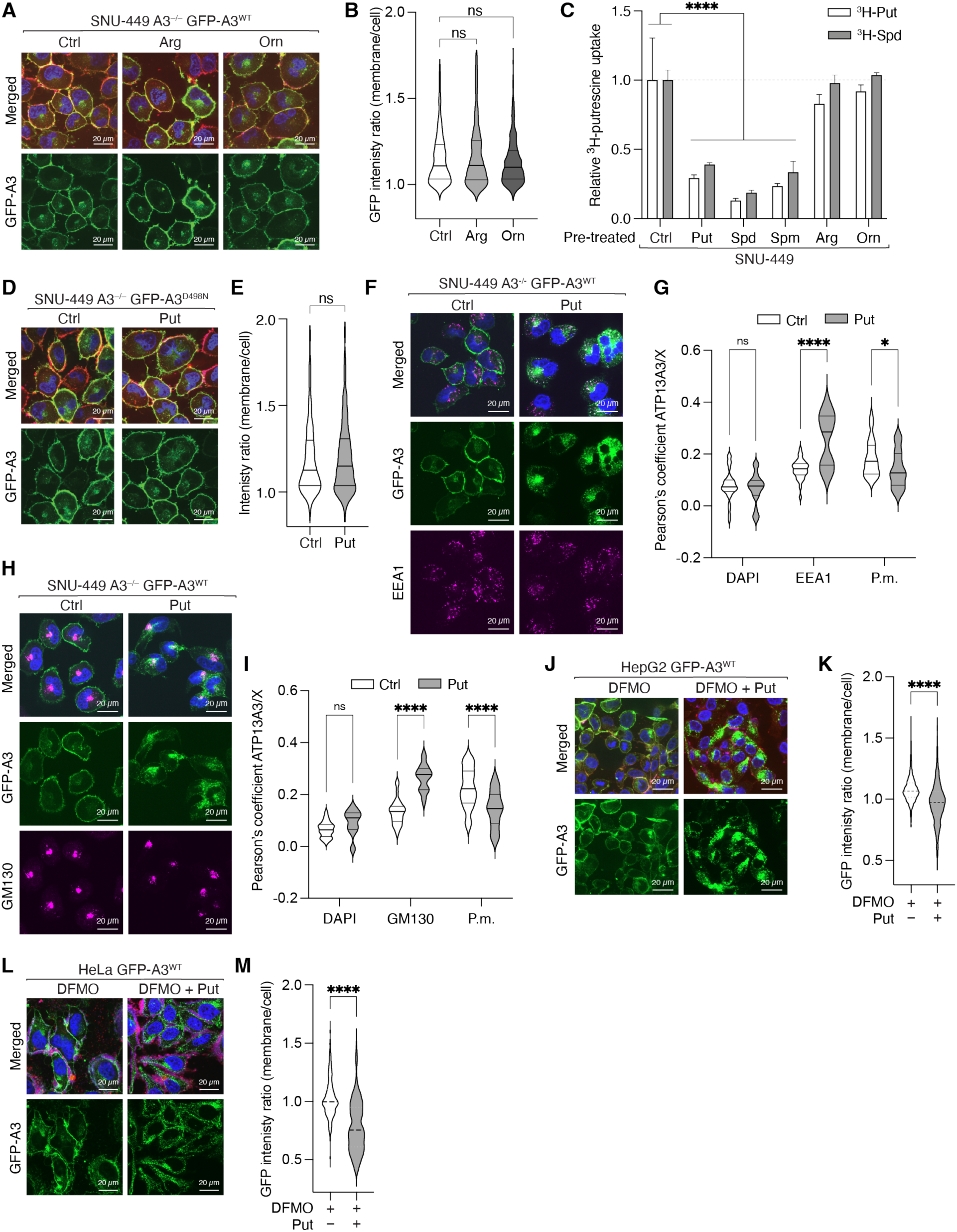
ATP13A3 changes its localization depending on intracellular polyamine levels. **(A** and **B)** Fluorescence microscopy of SNU-449 A3^-/-^ GFP-A3^WT^ cells treated with PBS, 10 mM arginine or 1 mM ornithine for 2 h. (**C**) Relative ^3^H-polyamine uptake into SNU-449 control cells treated with PBS, 10 µM putrescine, 10 µM spermidine, 10 µM spermine, 10 mM arginine, or 1 mM ornithine for 2 h. n=4. (**D** and **E)** Fluorescence microscopy of SNU-449 A3^-/-^ GFP-A3^D498N^ cells treated with 10 µM putrescine for 2 h. (**F** to **I**) Fluorescence microscopy and colocalization analysis of GFP-A3 with DAPI, plasma membrane marker, and EEA1 or GM130 in SNU-449 A3^-/-^GFP-A3^WT^ cells treated as in (D). (**J** and **K**) Fluorescence microscopy of HepG2 GFP-A3^WT^ cells depleted of polyamines (1 mM DFMO for 2 days) and treated with PBS or 10 µM putrescine for 6 h. (**L** and **M**) Fluorescence microscopy of HeLa GFP-A3^WT^ cells depleted of polyamines (1 mM DFMO for 2 days) and treated with PBS or 10 µM putrescine for 2 h. *p < 0.05, ****p < 0.0001 by one-way-ANOVA (B), two-way-ANOVA (C, G, I), or t-test (E, K, M).

**Fig. S4.**
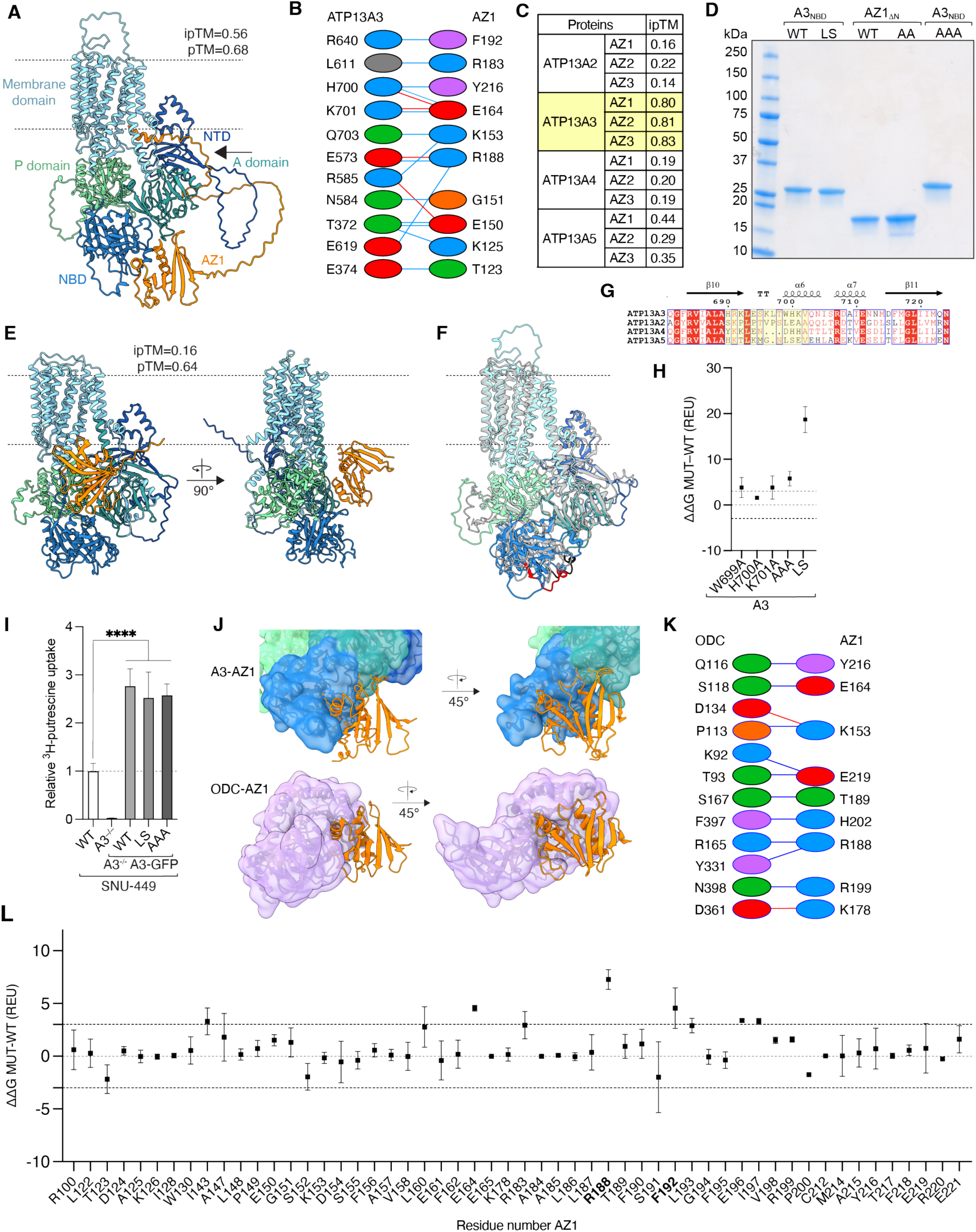
ATP13A3 interacts with AZ1 via its N-domain. **(A)** AlphaFold 3 prediction of the ATP13A3-AZ1 complex. Arrow indicates interaction of the AZ N-terminus with ATP13A3. (**B**) PDBsum analysis of the ATP13A3-AZ1_ΔN_ interaction interface. Residues are colored by physicochemical property: negatively charged (red), positively charged (blue), aromatic (purple), polar uncharged (green), and hydrophobic (orange). Hydrogen bonds and salt bridges between interfacial residues are depicted as blue and red lines, respectively. (**C**) AlphaFold 3 ipTM scores of all possible interactions between ATP13A2-5 and AZ1-3. (**D**) SDS-PAGE analysis of A3_NBD_ and AZ1_ΔN_ variants purified from *E. coli* and used for SEC-MALS experiments. (**E**) AlphaFold 3 prediction of the ATP13A2-AZ1 complex. (**F**) Superposition of AlphaFold 3 predicted ATP13A3 (colored) and ATP13A2 (gray) models. Loop swap region of ATP13A3 and corresponding region in ATP13A2 are colored in red and black, respectively. (**G**) Sequence alignment of a segment of the P5B-ATPase family NBDs. Loop swap region is highlighted in yellow. (**H**) *In silico* analysis of changes in binding energies between ATP13A3 variants and AZ1_ΔN_ using the Rosetta Interface Analyzer (RIA). Positive values indicate reduced binding energies. n=10 relaxation runs. (**I**) Relative ^3^H-putrescine uptake into SNU-449 control, A3^-/-^, A3^-/-^ GFP-A3^WT^, A3^-/-^ GFP-A3^LS^, and A3^-/-^ GFP-A3^AAA^ cells. n=4. (**J**) Comparison of the AlphaFold 3 predicted ATP13A3-AZ1 complex (top) and ODC-AZ1_95-221_ crystal structure (bottom; PDB: 4ZGY). ODC is colored in light purple, all other proteins are colored as in (A). (**K**) PDBsum analysis of the ODC-AZ1_ΔN_ interaction interface. Residues and interactions colored as in (B). (**L**) *In silico* alanine screening of the AZ1_ΔN_-ATP13A3 complex using the RIA. REU=Rosetta energy units. n=10 relaxation runs.

**Fig. S5.**
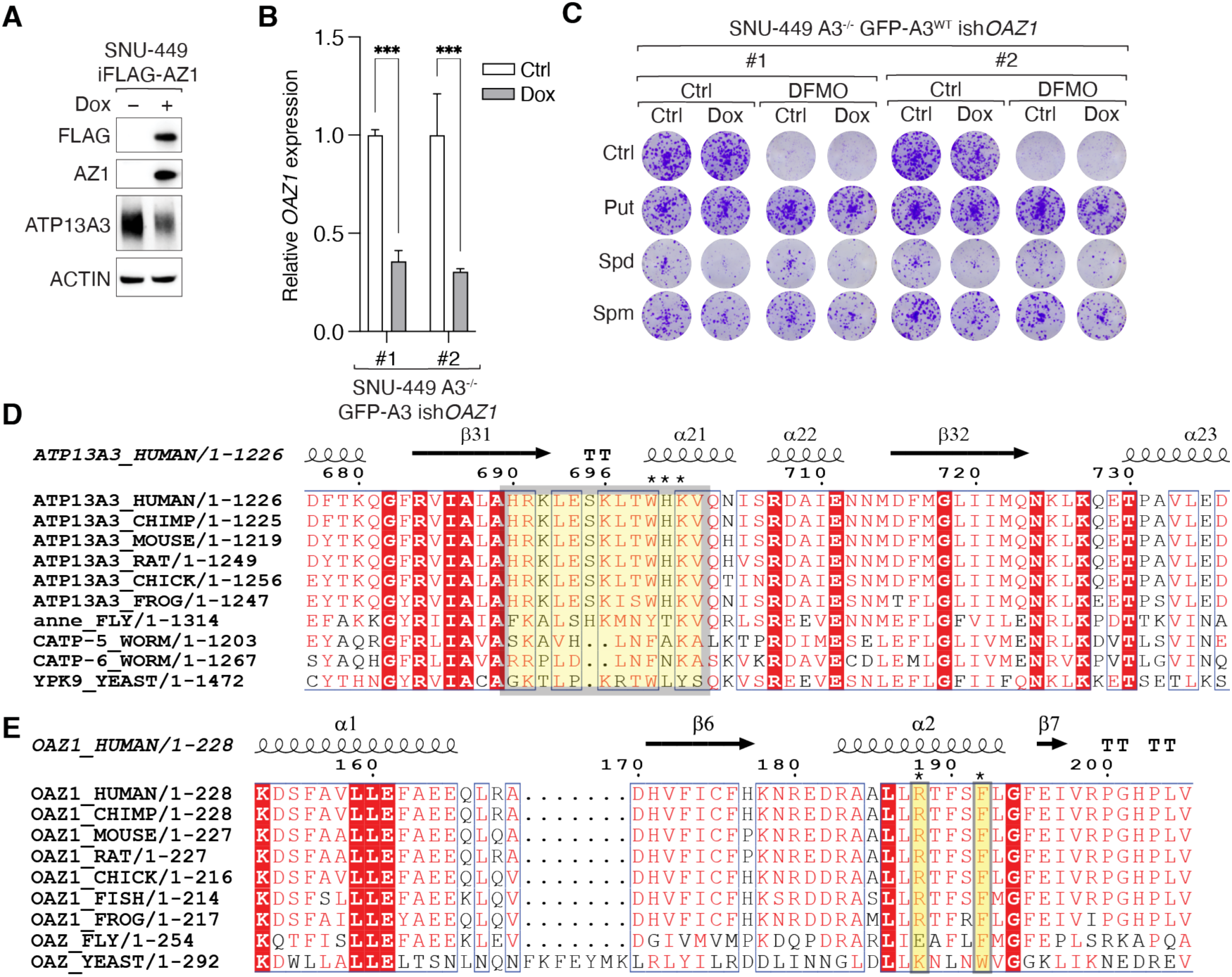
ATP13A3 and AZ1 interaction surface is necessary for polyamine homeostasis. (**A**) Immunoblots of FLAG, AZ1, and ATP13A3 in SNU-449 iFLAG-AZ1 cells. ACTIN serves as loading control. (**B**) mRNA levels of *OAZ1* upon siRNA-mediated *OAZ1* knockdown in SNU-449 cells. (**C**) Representative clonogenic growth assay of two SNU-449 A3^-/-^ GFP-A3^WT^ ish*OAZ1* clones in control or polyamine-depleted (1 mM DFMO for 2 days) conditions treated with DMSO or 1 µg/ml doxycycline and supplemented with 1 µM polyamines. (**D** and **E**) Multiple sequence alignments of a conserved region within ATP13A3 NBD or AZ1 from representative organisms. Residues critical for ATP13A3-AZ interaction are highlighted in yellow.

**Table S1.**
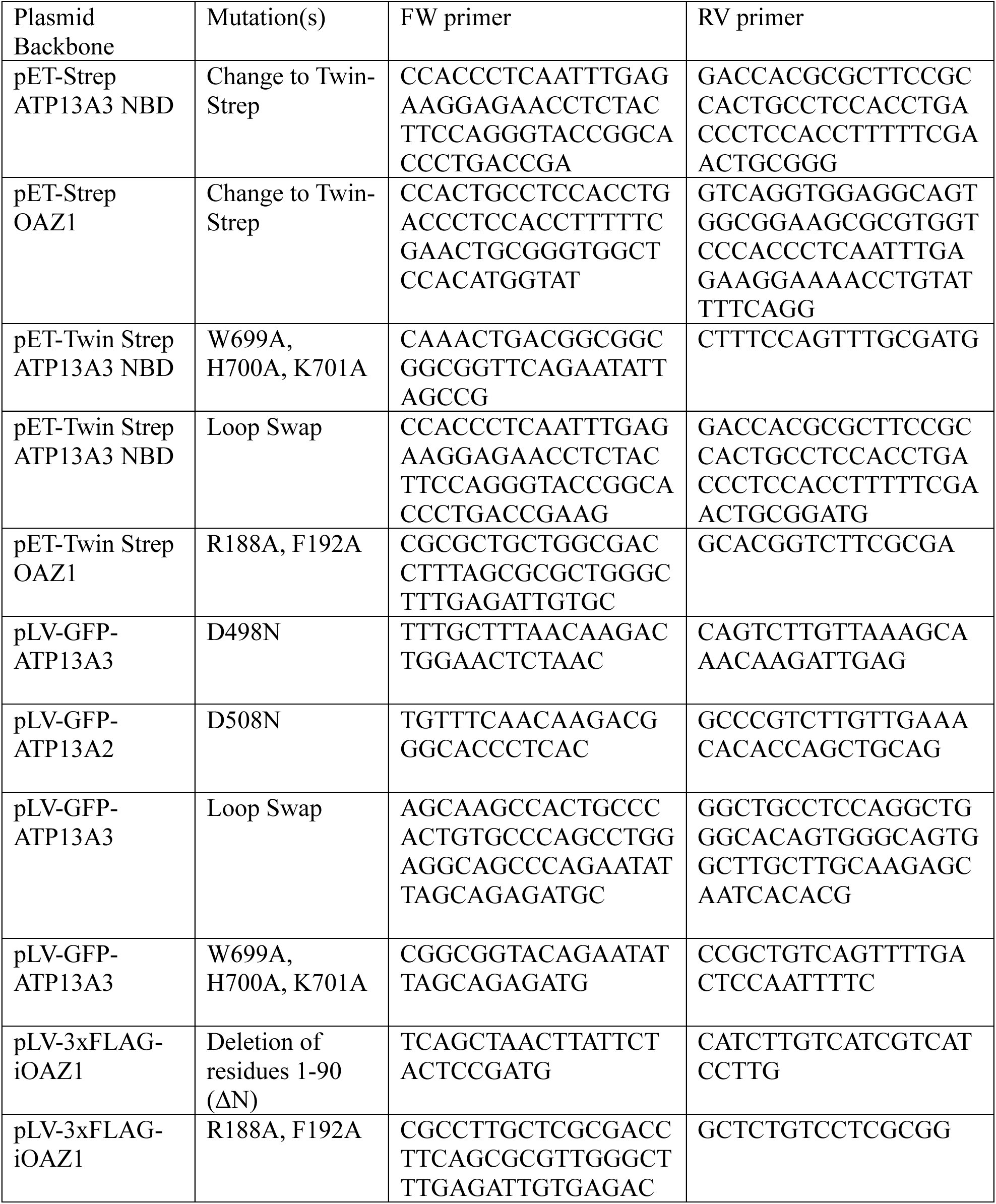
Primers used for mutagenesis of *ATP13A3* and *OAZ1* vectors.

